# Capturing site-to-site variability through Hierarchical Bayesian calibration of a process-based dynamic vegetation model

**DOI:** 10.1101/2021.04.28.441243

**Authors:** Istem Fer, Alexey Shiklomanov, Kimberly A. Novick, Christopher M. Gough, M. Altaf Arain, Jiquan Chen, Bailey Murphy, Ankur R. Desai, Michael C. Dietze

## Abstract

Process-based ecosystem models help us understand and predict ecosystem processes, but using them has long involved a difficult choice between performing data- and labor-intensive site-level calibrations or relying on general parameters that may not reflect local conditions. Hierarchical Bayesian (HB) calibration provides a third option that frees modelers from assuming model parameters to be completely generic or completely site-specific and allows a formal distinction between prediction at known calibration sites and “out-of-sample” prediction to new sites. Here, we compare calibrations of a process-based dynamic vegetation model to eddy-covariance data across 12 temperate deciduous Ameriflux sites fit using either site-specific, joint cross-site, or HB approaches. To be able to apply HB to computationally demanding process-based models we introduce a novel emulator-based HB calibration tool, which we make available through the PEcAn community cyberinfrastructure. Using these calibrations to make predictions at held-out tower sites, we show that the joint cross-site calibration is falsely over-confident because it neglects parameter variability across sites and therefore underestimates variance in parameter distributions. By showing which parameters show high site-to-site variability, HB calibration also formally gives us a structure that can detect which process representations are missing from the models and prioritize errors based on the magnitude of the associated uncertainty. For example, in our case-study, we were able to identify large site-to-site variability in the parameters related to the temperature responses of respiration and photosynthesis, associated with a lack of thermal acclimation and adaptation in the model. Moving forward, HB approaches present important new opportunities for statistical modeling of the spatiotemporal variability in modeled parameters and processes that yields both new insights and improved predictions.

## 1. Introduction

In times of unprecedented environmental change, process-based models are one of the most important tools at our disposal to understand and predict ecosystem function under global change. By encoding our current understanding about how processes work, these models often provide greater confidence when trying to understand current conditions, which are far from equilibrium and dominated by transients, as well as when making short-term ecological forecasts or long-term projections of environmental change (Dietze et al., 2018). However, it can be hard to apply these models effectively because the parameters in such models are empirical coefficients that need to be calibrated to data, which can vary in space and time (Zaehle et al., 2005; Buotte et al., 2021). This presents a conundrum of how to calibrate with data that are collected over different sites or other observational units, be it stands, individuals, watersheds, lakes depending on the scales that models operate and the scales that data are collected.

Users are thus often presented with a choice between using general model parameters or site-specific parameters. General parameters may reflect uncalibrated “defaults”, or may result from performing a multisite “joint” calibration against all the data from all the sites assuming no site-to-site variability (Minunno et al., 2019, Mäkelä et al., 2020). But in either case such parameter sets often have mixed success at capturing the dynamics at any particular site. Site-specific calibrations, on the other hand, assume all sites to be completely independent but generally do a much better job of capturing site-level dynamics, although can be both labor- and data-intensive to generate (van Oijen, 2017) and limit potential for upscaling. Similar to general calibrations, site-specific calibrations also have had mixed success in their ability to make predictions at new sites (Post et al., 2017). Furthermore, while ecologists have only recently begun propagating uncertainties in process-based models, both the general and site-specific approaches are likely to not just be ‘wrong’ when applied to new sites, but to be over-confidently wrong or unquantifiably uncertain, with confidence intervals that do not reflect the true uncertainties associated with extrapolating calibrations to new sites. This out-of-sample overconfidence is occurring precisely because neither approach has a way of quantifying or propagating the uncertainties associated with the spatiotemporal variability in model parameters. Finally, the presence of spatiotemporal variability in model parameters itself implies that models are missing important interactions and processes altogether (Medlyn et al., 2015).

Hierarchical Bayesian (HB) approaches to calibration provide a third option in the continuum between these two approaches (Fig 1) that allows us to quantify and partition the scales of unexplained uncertainties (what statisticians call random effects, or in our specific case; site effects), and frees the modelers from assuming model parameters to be completely generic or completely site-specific (van Oijen, 2017). The hierarchical calibration framework has two levels: at the site-level, parameters are fitted to each site, then at the hierarchical level, sites are modeled to be distributed around an across-site (global) mean with an across-site (global) variance. After calibration, when a new prediction is made at one of the sites that went into the hierarchical calibration (in-sample prediction), the site-level parameters for that site will be used. When a new prediction will be made at a new site that was not part of the hierarchical calibration (out-of-sample prediction), global level parameters will be used. HB calibration also borrows strength from sites with more data and constrains models at under-sampled sites, which is a common scenario in ecology (Clark, 2005). If the sites are very different (large site-to-site variability), the across-site model would provide very little constraint on parameters. If the sites are very similar (little or no site-to-site variability) the across-site model would behave very much like the joint calibration. However, in either case this structure is formally informed by the data rather than making *a priori* assumptions. This structure also allows us to constrain parameters at one site based on observations at another when there are high across-site correlations among parameters.

**Fig 1.**
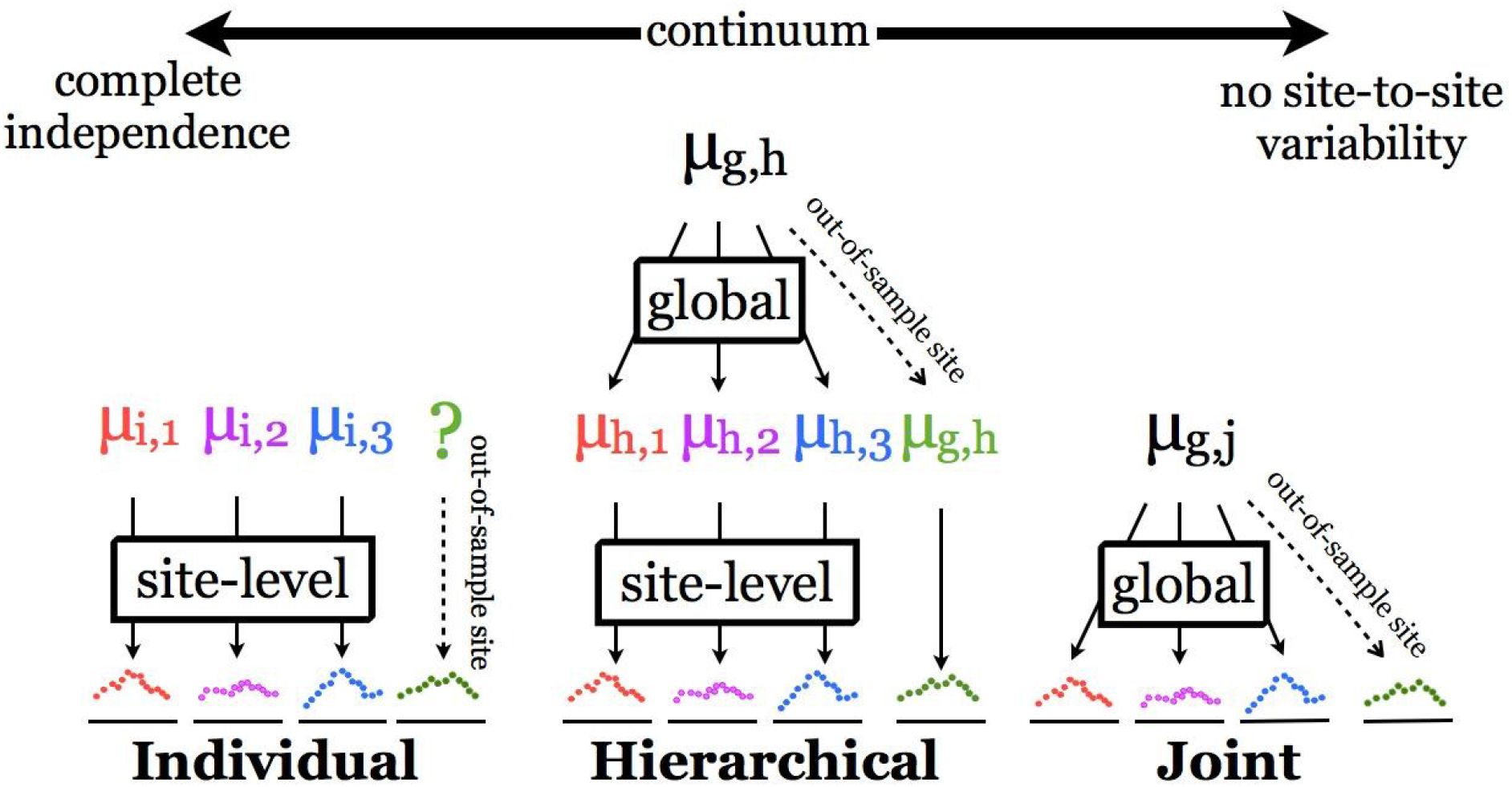
Conceptual figure showing that Hierarchical Bayesian calibration is on the continuum of individuallevel calibration where we assume complete independence and joint calibration where we ignore all among site variability (after Dietze, 2017). μ_i,1/2/3_, μ_h,1/2/3_ : site-level parameters under individual and hierarchical calibration, respectively. μ_g,h/j_: global parameters under hierarchical and joint calibration, respectively.

HB not only provides improved predictions, but also formally gives us a structure that will help us better understand this inter-site variability moving forward (Dietze, 2017). By looking at which parameters show the highest site-to-site variability, we can detect both known and novel missing key processes. For example, one of the known missing and influential processes from dynamic vegetation models is the temperature acclimation of photosynthesis (Dietze, 2014; Lombardozzi et al., 2015). In a HB calibration, this might be detected as optimum photosynthesis parameters showing high site-to-site variability, as the calibration would try to compensate for this missing photosynthesis process in the model by suggesting different values for different sites. However, the reason why some of these parameters show high site-to-site variability may not always be known, and may point to new questions that we did not think to ask until we applied the HB framework. In such cases, we can generate hypotheses and perform simple post-calibration analyses where we can test which predictors can explain this site-to-site variability (Fer, 2020). Furthermore, by looking at across-site parameter correlations we can detect model structural errors (assuming the trade-offs in across-site parameters are not biologically meaningful). Overall, this framework gives us a way of detecting whether a process needs to be added to a model without adding it and seeing if the model gets better.

HB approach has recently gained attention for the calibration of process-based models (Thomas et al., 2017; Susiluoto et al., 2018; Tian et al., 2020), however, hierarchical process-based model calibrations are still very limited. One reason why a HB approach may not have yet been widely adopted by the process-based modeling community is that Bayesian calibration is already computationally costly for process-based models (Fer et al., 2018). Recently, statistical and computational solutions for calibration of process-based models have gained attention, including Bayesian model emulation (Fer et al., 2018), which could help overcome this hurdle. Here we build upon our previous emulator-based Bayesian calibration tools to develop a generalized, emulator-based multi-site HB calibration tool. In addition, this tool is coupled with automated pipelines within the Predictive Ecosystem Analyzer (PEcAn), a community cyberinfrastructure tool, that can handle the data processing and execution of large numbers of model runs across a large number of sites (LeBauer et al., 2013; Fer et al., 2021).

In this study, we assimilated data from the Ameriflux network to calibrate a process-based terrestrial ecosystem model, Simplified Photosynthesis and Evapotranspiration (SIPNET), using the HB approach and present insights gained from it in comparison to both individual site-level and “joint” across-site simple Bayesian (SB, as referred by Clark, 2005) approaches (Fig 1). We specifically seek to test the following hypotheses about in-sample calibration and prediction (H1 and H2), out-of sample prediction (H3) and uncertainty partitioning (H4):

(H1) Model predictions will be falsely over-confident after joint fitting because it neglects across-site parameter variability and estimates a single parameter vector for all sites. In other words, uncertainties in parameter posterior estimates and post-calibration model ensemble output will be narrower but model-data agreement will be worse (higher deviance).
(H2) Model-data comparison will show the best agreements (lowest deviance) after the site-level calibrations. Sites-level parameter estimates will gain additional constraints from HB calibration, and HB site-level post-calibration model ensemble agreement with data will be better (lower deviance) than SB site-level post-calibration ensemble.
(H3) Out-of-sample model predictions using parameters fitted through joint calibration will again be overconfident: i.e. predictive intervals will be more precise (narrower width) but will often exclude data at out-of-sample sites (lower coverage). Model predictions using the HB calibration will be more uncertain (reflecting the true uncertainties) but more accurate (better model assessment scores) at out-of-sample sites.
(H4) Across-site parameter variability will be higher than parameter uncertainty.

## 2. Methods

### 2.1 Study sites and model

The Ameriflux network provides long-term standardized measurements across a wide range of climatic and geographic space (Novick et al., 2018). Here we limited our study system to Eastern U.S. Temperate Hardwoods (categorized as ‘Deciduous Broadleaf Forest’ -DBF- in Ameriflux) to avoid inherent parameter differences from use of the Plant Functional Type formalism in ecosystem models and minimizing bioclimatic variance as an additional potential source of parametric uncertainty. The 12 Ameriflux DBF sites with available data, in terms of both meteorological drivers and data to assimilate (CA-TPD, Arain, 2018; CA-Oas, Black, 2016; US-Bar, Richardson & Hollinger, 2019; US-ChR, Meyers, 2016; US-Dk2, Oishi et al., 2018; US-Ha1, Munger, 2020; US-MMS, Novick & Phillips, 2020; US-MOz; Wood & Gu, 2019; US-Oho, Chen et al., 2019; US-Slt, Clark, 2016; US-UMB, Gough et al., 2021; US-WCr, Desai, 2021), are shown in Fig. S1 on their climatic and geographic space, and are also listed in Table S1 with the longest uninterrupted (continuous) data availability range given in the “Years used” column. Differences in number of active tower years among sites provides an opportunity to demonstrate the ability of HB calibration to borrow strength across sites that vary in information availability. Among available Ameriflux data, we use Net Ecosystem Exchange (NEE) and Latent Heat (LE) as model constraints on carbon and water dynamics, respectively.

To illustrate our Hierarchical Bayes (HB) calibration, we used the Simplified Photosynthesis and Evapotranspiration (SIPNET) model (https://github.com/PecanProject/sipnet), which we ran through the Predictive Ecosystem Analyzer (PEcAn, LeBauer et al., 2013; https://github.com/PecanProject/pecan) ecological informatics toolbox. SIPNET is ideal for our study as it was developed to facilitate model comparisons to eddy-covariance data (Braswell et al., 2005; Sacks et al., 2006).

For all of our calibration and validation runs we ran SIPNET’s temperate deciduous PFT for the given periods (ranging from 3 to 14 years of data, also see Table S1, “Years used” column) using Ameriflux tower meteorological data as model drivers. Meteorological drivers were gap-filled using the marginal distribution sampling approach implemented in the REddyProc package described in Wutzler et al. (2018). NEE and LE data were not gap-filled as this is not required for calibration and doing so would have artificially inflated sample sizes and resulted in a model-model comparison. SIPNET also requires initial conditions for its leaf, wood, and soil C pools (Table S2), which we extracted from the Ameriflux BADM (Biological, Ancillary, Disturbance and Metadata) files, or from reference papers listed in the BADM files, when possible. For the remaining sites, we used other papers in the literature, personal communication, and expert opinion. The initial conditions used at each site are given in Table S3, and sources used in their compilation listed in Table S4.

### 2.2 Parameter selection

To reduce the dimensionality of the calibration we screened SIPNET’s soil decomposition and plant physiology parameters (40 in total) to determine which parameters contribute most to the uncertainty in the NEE and LE predictions of the model, and thus could be constrained by NEE and LE data in calibration. We used a global uncertainty analysis approach to determine the contribution of different parameters to the output uncertainty. First, we generated 250 ensemble runs per site by uniformly sampling prior parameter distributions. Next, we fitted separate univariate linear regressions between all 40 parameter samples and outputs. Then, we estimated the relative importance of each parameter using the respective R^2^ values of each linear regression (Dietze, 2017). Full sensitivity analysis results are available in Fig. S2. In the end, we selected the 5 most influential parameters (Table 1) which combined explained 60% and 53% of model LE and NEE output variability on average (for each site they explain a slightly different amount), respectively.

**Table 1.** Top 5 SIPNET parameters that contributed most to the model output (NEE and LE) uncertainty.

| Parameter | Definition | Units | Prior |
| --- | --- | --- | --- |
| wueConst | Water use efficiency constant | unitless | Unif[0.001, 50] |
| psnTOpt | optimum photosynthesis temperature at which net photosynthesis occurs | °C | Gamma[5, 0.2] |
| rdConst | scalar determining canopy aerodynamic resistance | unitless | Unif[1, 1500] |
| som_respiration_rate | Soil organic matter respiration rate coefficient | day-1 | Unif[0.001, 0.2] |
| leafGrowth | Carbon allocated to leaf flush at the start of growing season | gC m-2 | Weibull[2, 150] |

### 2.3 Model calibration experiments

In this study, we compare the performance of three calibration approaches (Fig. 1): i) individual site-level, ii) joint across-site, and iii) hierarchical. Note that both individual and hierarchical approaches produce site-level calibrations. Therefore, we specify the site-level calibrations in the individual calibration framework as simple Bayesian (SB) site-level, and in the hierarchical framework as HB site-level (in-sample). Likewise, both joint and hierarchical calibration produces “global” across-site parameters which are referred as joint-global and hierarchical-global, respectively (Table 2). All results are reported for the 12 DBF sites.

**Table 2.** Ensemble experiments designed in this study. Each experiment used an ensemble size of 250 runs at each of 12 sites. Last column indicates whether all parameters were varied in the ensemble run (Yes) or just the five targeted parameters (No).

| Calibration approach | Description | All pars |
| --- | --- | --- |
| Pre-PDA ensembles | Model is run with PEcAn meta-analysis posteriors (PDA-priors). | Yes |
| SB site-level | Model is run in ensemble mode with individual site-level PDA posteriors at each site. | Yes |
| HB site-level (in-sample) | All 12 sites are included in hierarchical calibration. Then, the model is run in ensemble mode with hierarchical site-level PDA posteriors at each site. | Yes |
| Joint (in-sample) | All 12 sites are included in joint calibration. Then, the model is run in ensemble mode with joint global posteriors at each site. | Yes |
| HB / LOO-CV (out-of-sample) | HB calibration is re-run by leaving one site out at a time. Then, the model is run in ensemble mode and once with MAP using hierarchical global posteriors at the site that is left out. | Yes |
| Joint / LOO-CV (out-of-sample) | Joint calibration is re-run by leaving one site out at a time. Then, the model is run in ensemble mode and once with MAP using joint global posteriors at the site that is left out. | Yes |
| HB site-level (5 parameters) | Same calibration as the HB site-level (in-sample) above. Then, the model is run in ensemble mode with hierarchical site-level PDA posteriors at each site, varying only the 5 SIPNET parameters. | No |
| HB global (5 parameters) | Same as HB / LOO-CV (out-of-sample) above. Then, the model is run in ensemble mode using hierarchical global posteriors at the site that is left out, varying only the 5 SIPNET parameters. | No |

PDA: Parameter Data Assimilation, LOO-CV: leave-one-out cross validation, MAP: Maximum a posteriori estimation

One of our hypotheses (H3) suggests that model predictions with hierarchical global posteriors will capture data at out-of-sample sites better than the model predictions using joint global parameters. To test this, we used leave-one-out cross validation (LOO-CV, Gelman et al., 2013; Susiluoto et al., 2018) where we repeated the joint and HB calibration by leaving one site out at a time, and using the global parameter estimates for prediction at the out-of-sample sites. LOO-CV was not performed for the independent site-level SB fits because, from a Bayesian perspective, this approach does not provide a basis for out-of-sample prediction.

Table 2 lists the calibration analyses conducted at each site. Each experiment used an ensemble size of 250 runs at each of 12 sites (and two additional MAP runs for LOO-CV experiments), with a total of 3000 runs (3024 for LOO-CV) per site. Pre- and post-calibration performance was compared using (normalized) deviance, root mean square error (*RMSE*), bias, correlation coefficient (*R*), slope of the regression (*R^2^*), coverage, and width of prediction intervals as metrics, in addition to visual comparisons.

### 2.4 Emulator protocol

In all three calibration frameworks (individual, hierarchical, joint), we used the emulator approach that is implemented in PEcAn and explained in detail by Fer et al. (2018). Emulators are statistical models that are used in place of the full model and are typically much faster to run. Although SIPNET is decently fast for a process-model (hence, could be run in a Markov Chain Monte Carlo -MCMC- loop sequentially for 10^4^-10^7^ times), Fer et al. (2018) showed substantial gain in terms of computation time using the emulator approach with calibration performances that are comparable to using the full model in the MCMC. These gains are going to be even more pronounced for the HB case, where the use of emulators allows us to avoid rerunning the process-based model when new data and/or new sites are added or when more complex statistical models are explored at the hierarchical level (e.g. additional spatiotemporal structure or cross-site covariates, see *Discussion*). Moreover, this study aims to demonstrate the strength of HB calibration, and promote its adoption in the process-based modeling studies where the computational costs of Bayesian calibration algorithms have been a hurdle. Therefore, in this study we used the emulator approach which can be readily extended to more complex models than SIPNET.

The overall calibration scheme closely follows Fer et al. (2018): Calibration priors for model parameters were generated by PEcAn’s trait data meta-analysis module, which performs a univariate hierarchical Bayesian meta-analysis while accounting for random site effects (LeBauer et al., 2013; Raczka et al., 2018). Flux data were u* filtered, corrected for autocorrelation and residuals were modeled by a heteroskedastic Laplacian error distribution following Richardson et al. (2006). Gaussian Process (GP) models were used as the emulator following the basic algorithm:

1. Propose *N_knots_* parameter vectors
2. Run model (in parallel)
3. Calculate model-data errors
4. Fit an emulator for each site (*S*), built on their likelihood surfaces (*GP_S_*)
  4a. Optional: iterative refinement of error surface
5. Perform calibration using *GP_S_*.

Each site-level emulator calibration is run in iterative rounds with adaptive sampling following Fer et al. (2018), with the exception that we changed the total number of proposed knots per round in this study to *px30, p* being the number of parameters. This adaptive sampling achieves an effect analogous to a nested grid, providing a higher resolution near the center of the posterior distribution and sparse sampling in parts of parameter space with low probability. Fer et al. (2018) showed that gain in terms of reduction in emulator approximation error starts to be heavily traded off with the compute time of the emulator itself after certain numbers of knots. Accordingly, we repeated the iterations until such threshold was reached, and stopped after the 4^th^ round (# of rounds x parameters x knots = 4 x 5 x 30 = 600 knots in total) as the 5^th^ round posterior distributions were not dramatically improved in comparison (not shown). This number is also in agreement with the scaling experiment in Fer et al. (2018) where they showed the rate of improvement in model data agreement has not substantially changed as proposed number of knots reached 480 for 6 parameters (Fig 7b therein).

### 2.5 Calibration scheme

Algorithms for all three workflows are provided in Fig 2. Site-level emulators are built over the same set of knots at all 12 sites. This was done to minimize the differences due to GP interpolation uncertainty in the likelihoods returned by the emulators at different sites. This way, each emulator at each site passes exactly through the same design points where the model has been run. SB site-level calibration is performed by proposing new parameter vectors for each site and by estimating individual posteriors through each *GP_S_* independently in the MCMC (Fer et al., 2018). In joint calibration MCMC, a proposed parameter vector is accepted or rejected according to the estimated posterior values using all *GP_S_* jointly (i.e. using the product of the Likelihoods across all sites). In the hierarchical MCMC, site-level parameter vectors, μ_S_, are accepted or rejected at the site-level using the across-site mean, μ_g_, and precision, τ_g_, as priors. The across-site mean and precision matrix are updated simultaneously through Gibbs sampling to further improve the computational efficiency of our workflow.

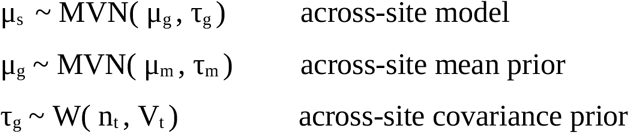

**Fig 2.**
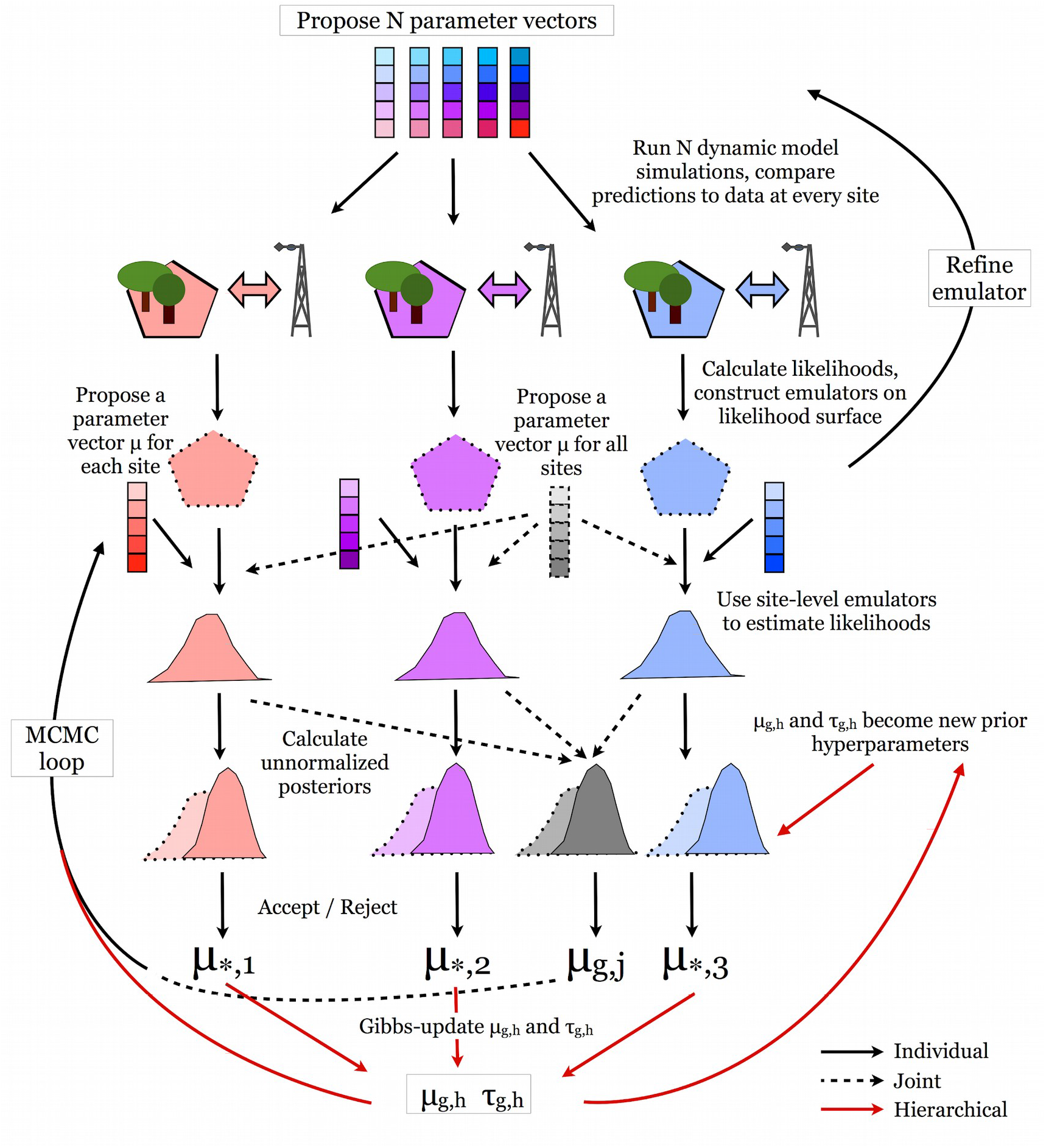
Workflows for all three calibration schemes. First (top), emulators are constructed on likelihood surfaces as explained in Fer et al. (2018). Then (bottom, Markov Chain Monte Carlo -MCMC- loop), new parameter vectors are proposed, unnormalized posteriors are calculated, proposed parameter vectors are accepted/rejected. In the individual calibration (solid, black lines), sites are fitted independently to obtain posteriors of μ_i,1/2/3_ (* is a wildcard for *i* in individual, for *h* in hierarchical site-level calibration, also see Fig 1). In the hierarchical calibration (solid, black lines followed by solid, red lines), global posteriors μ_g,h_ are fitted simultaneously using an additional Gibbs update step, and used as the new prior hyperparameters in the next iteration of MCMC to obtain site-level hierarchical posteriors, μ_h, 1/2/3_. In the joint calibration (black, dashed lines), sites are fitted together (i.e. a new proposed parameter vector is accepted/rejected depending on joint likelihood estimates from all sites) to obtain joint global posterior μ_g,j_.

For Gibbs updating, we chose the hyperpriors for the across-site mean, μ_g_, and precision, τ_g_, to be multivariate Normal and Wishart for conjugacy. The prior parameters for the across-site mean were set based on the posteriors from the trait meta-analysis, analogous to the SB priors (Table 1). The hyperparameters for the Wishart across-site precision prior were set to uninformative values of *n_t_=6=p+1, p* being the number of parameters, and *V_t_=∑_m_/n_t_, ∑_m_* being the covariance matrix, i.e. inverse of hyperparameter *τ_m_*.

### 2.6 Analysis of the random effects

The hierarchical model partitions the overall variability into two terms, one describing the within-site parameter uncertainty and the other describing across-site variability. To aid interpretation we first decompose the across-site (global) covariance matrix (Σ = D·R·D) into an across-site parameter correlation matrix (R) and a vector of standard deviations (D). We then converted the standard deviations to unitless coefficients of variation to determine which parameters show the most variation and which are conserved among sites. Next, to investigate what factors might explain site-to-site parameter variability we performed a series of exploratory correlation analysis between the maximum a posteriori (MAP) estimations for all parameters from all sites and a set of candidate variables. Specifically the following variables are proposed to correlate with site-to-site variability:

- Mean annual air temperature, precipitation, photosynthetically active radiation, vapor pressure, wind speed
- Initial litter, wood, soil C pools, and their total
- Precalculated observation error (Laplace) parameters and sample sizes

### 2.7 Analysis of uncertainty and variability

Finally, we performed an uncertainty analysis to partition the relative contribution of within versus across-site variability in model predictive uncertainty. To do so we propagated both the parameter uncertainty encapsulated by the site-level parameter posterior distributions, and parameter variability encapsulated by the global parameter posterior distributions into model predictions (Table 2, last two experiments). We then compared the variance of this prediction to an ensemble that only considered site-level parameter uncertainty. In these ensemble runs, not to obscure the analysis with the remaining uncertainties in the non-targeted parameters, only the five parameters that were targeted in the calibration were varied while the rest of the parameters were kept at their prior mean.

## 3. Results

### 3.1 Comparison of parameter posterior distributions

To assess the impact of calibration approach on parameter means and uncertainties we report the marginal posterior median and 95% ranges at each site for each calibration approach (Fig 3). As expected, the joint calibration global posteriors were the narrowest for all parameters as this approach requires estimating the smallest number of parameters (i.e. one parameter vector for all sites). The HB global posteriors, which describe the across-site means, were typically widest, but still more informed than the priors. This is also expected as the width of these interval estimates is primarily controlled by the number of sites, not the amount of data available within a site, and we only used 12 sites in this analysis. In general, the joint and HB global means were very similar for all parameters, with the notable exception being the aerodynamic resistance (rdConst) which was higher in the joint calibration and the optimum photosynthesis temperature (psnTOpt) which was lower in the joint calibration. More than half of the sites with less constrained SB site-level posteriors, such as US-ChR, US-Bar, US-Dk2, US-Oho, US-Slt, US-MMS and US-UMB had narrower HB site-level (in-sample) posteriors, suggesting these sites were being informed by other sites in the hierarchical calibration scheme (also see Fig 6). Density plots for each marginal posterior are also shown in Fig S3.

**Fig 3.**
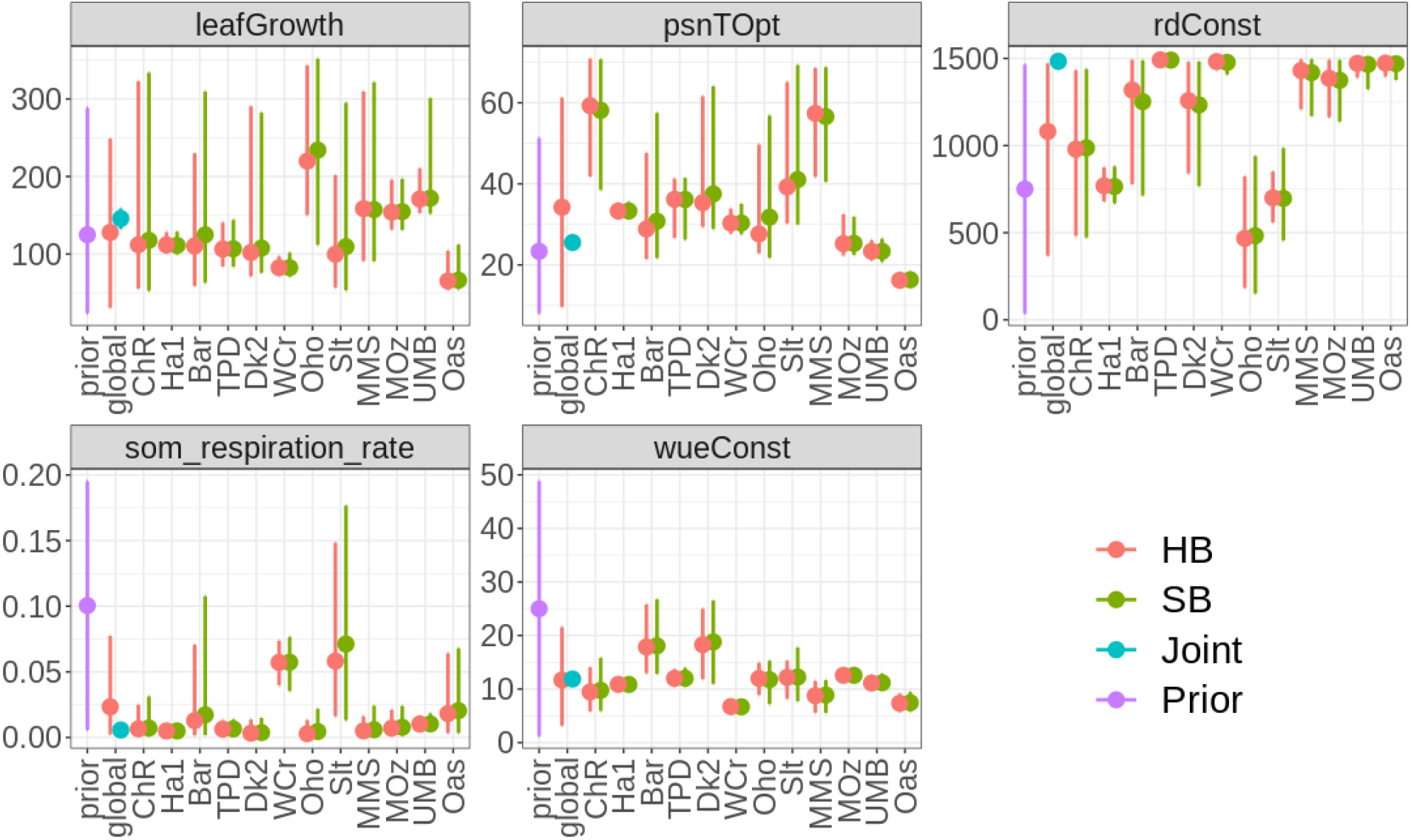
Comparison of marginal posterior 95% CI ranges after SB site-level, HB (in-sample) and joint (in-sample) calibration. All posteriors were more informed than the priors (purple bars). Joint calibration posteriors (blue bars) were the narrowest for all parameters. HB site-level (insample) marginal posterior ranges (red bars) were typically narrower than SB site-level posteriors (green bars) but not for all sites.

### 3.2 Analysis of random effects

Decomposing the across-site (global) covariance matrix showed that soil respiration parameter has the highest site-to-site variability (Fig 4, right panel). This is in contrast with the relatively modest variation apparent in Fig 3, especially in comparison to the prior range, but reflects the consistently low values of the posterior parameter estimates, causing the relative variability to be high. Across-site parameter correlations were generally modest; the parameters that showed the highest site-to-site (negative) correlations with each other (Fig 4, left panel) were the leaf growth and the canopy aerodynamic resistance parameter with −0.3, and leaf growth and the soil respiration parameter with −0.29.

**Fig 4.**
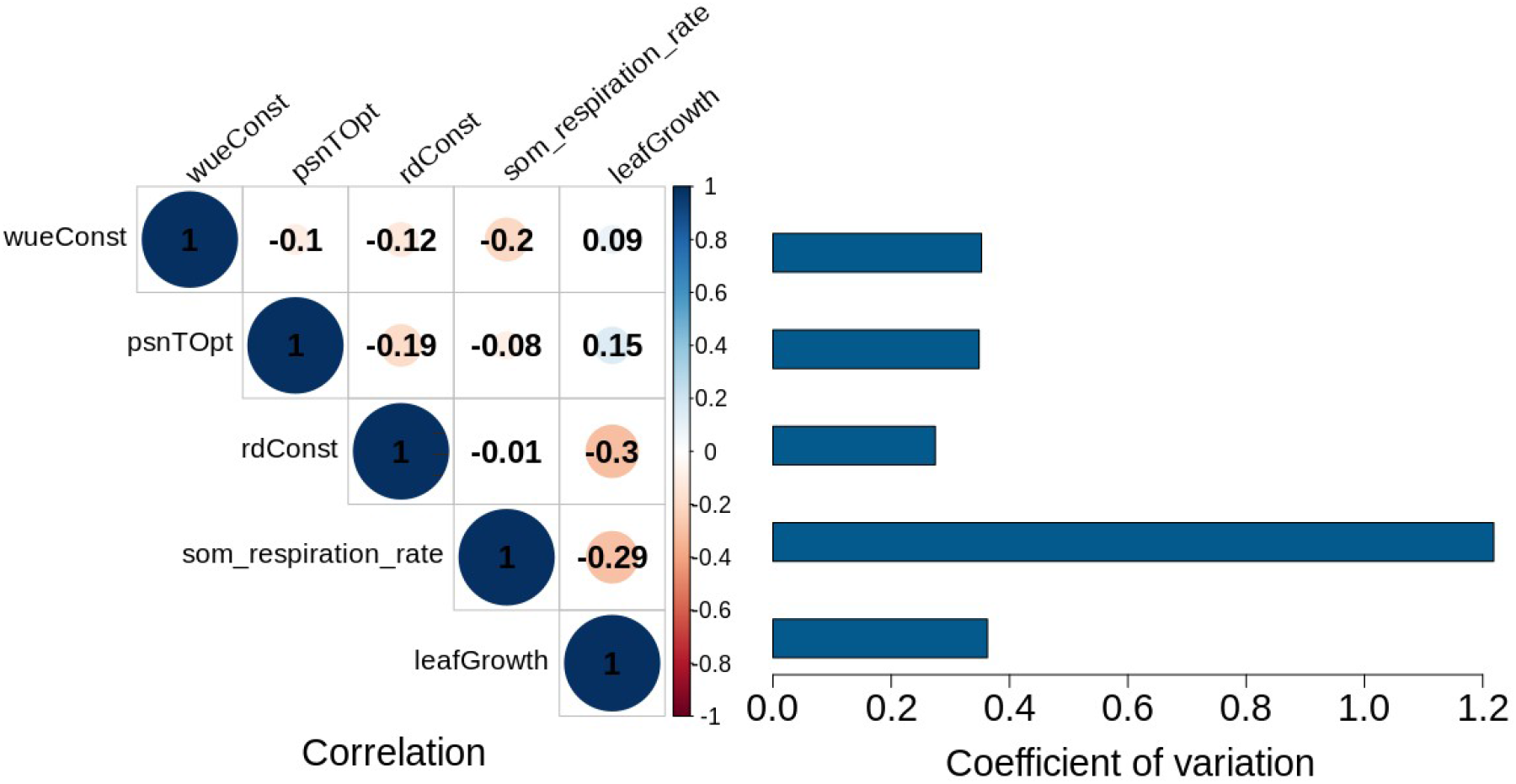
(Left panel) Across-site parameter correlations: Correlations among parameters allow us to constrain one parameter at one site based on observations at another. (Right panel) Across-site parameter standard deviations converted to unitless coefficients of variation: Parameters showing high site-to-site variability might imply missing processes in the model.

The exploratory correlation analysis (Fig 5) showed relatively high correlations between potential explanatory variables and across-site parameters: mean annual temperature, vapor pressure and precipitation showed positive correlation with optimum photosynthesis parameters (0.62 and 0.58 respectively, *p < 0.05*), initial soil carbon and total carbon showed negative correlation with soil respiration rate parameters (−0.53 and −0.66 respectively, *p < 0.05*), PAR showed negative correlation with the leaf growth parameters (−0.62, *p < 0.05*), wind speed showed positive correlation with the canopy aerodynamic resistance parameters (0.55, *p < 0.05*), wood carbon mass showed positive correlation with water use efficiency constant (0.67, *p < 0.05*). The precalculated parameters (intercept and negative slopes, Fig. S5) of the heteroskedastic Laplace showed high correlations with water use efficiency constant and optimum photosynthesis parameters (Fig 5).

**Fig 5.**
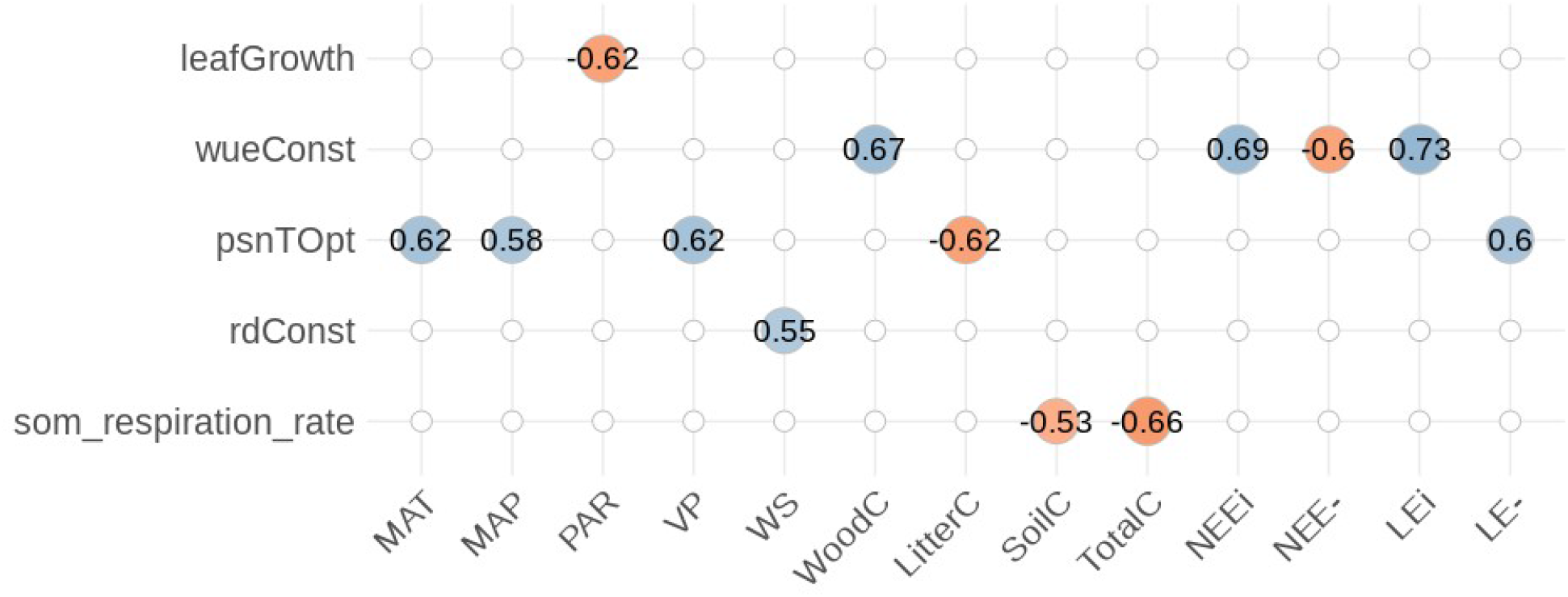
Exploratory correlation analysis results showing which factors may be explaining site-to-site parameter variability. Only variables that showed significant correlation (filled circles, p < 0.05) at least with one parameter are presented. MAT: mean annual temperature, MAP: mean annual precipitation, PAR: photosynthetically active radiation, VP: Vapor pressure, WS: wind speed, WoodC: wood carbon, LitterC: litter carbon, SoilC:soil carbon, TC: total (wood+litter+soil) carbon, NEE and LE i/n: Laplace parameters intercept (i), and negative (n) slopes of Net Ecosystem Exchange and Latent Heat, respectively.

### 3.3 Comparisons of calibration approaches at in-sample sites

(H1) Model predictions will be falsely over-confident after joint fitting because it neglects across-site parameter variability and estimates a single parameter vector for all sites. In other words, uncertainties in parameter posterior estimates and post-calibration model ensemble output will be narrower but model-data agreement will be worse (higher deviance).
(H2) Model-data comparison will show the best agreements (lowest deviance) after the site-level calibrations. Sites-level parameter estimates will gain additional constraints from HB calibration, and HB site-level post-calibration model ensemble agreement with data will be better (lower deviance) than SB site-level post-calibration ensemble.

Comparison of pre- and post-calibration fits supported the first two hypotheses (Fig 6). While all three calibration approaches showed improvement over the pre-PDA (Fig S4), the joint calibration usually had the largest deviance despite being more confident. Site-level HB deviances were typically as good or better than site-level SB calibration performance. This demonstrates that even when there are large amounts of data available at a site, in-sample (known site) prediction can be improved through HB calibration by leveraging the strength of additional information from across sites.

**Fig 6.**
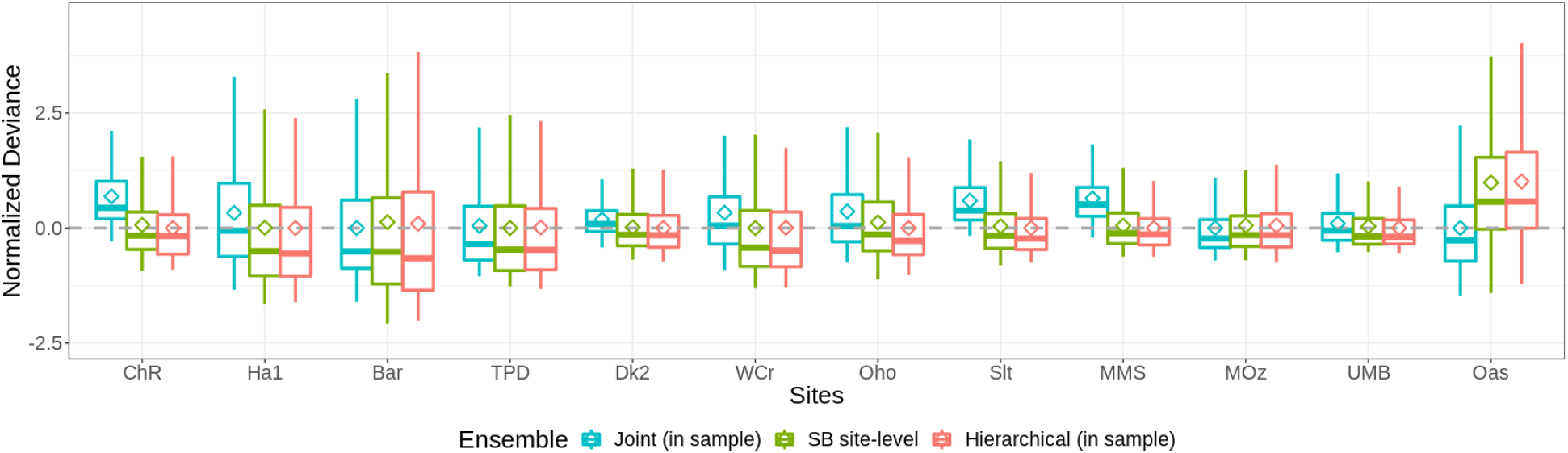
Deviance results for ensemble runs (ensemble size 250) with different calibration approaches listed in Table 2. The lower the deviance the better. The lower and upper hinges correspond to the first and third quartiles (the 25th and 75th percentiles) and the diamond shapes represent the mean. The deviance values are first normalized with respect to the sample size at each site, then the lowest ensemble mean among the calibration experiments was subtracted from all values which resulted in calibration with lowest mean appearing on the (dashed) zero line. Sites are ordered with respect to their data availability from left to right (less to more).

### 3.4 Comparisons of calibration approaches at out-of-sample sites

(H3) Out-of-sample model predictions using parameters fitted through joint calibration will again be overconfident: i.e. predictive intervals will be more precise (narrower width) but will often exclude data at out-of-sample sites (lower coverage). Model predictions using the HB calibration will be less precise (more uncertain) but more accurate (better model assessment scores) at out-of-sample sites.

The leave-one-out cross validation (LOO-CV) tests with joint and hierarchical global parameters at the out-of-sample sites partly supported the third hypothesis. Figure 7 reports performance metrics for the MAP run and the ensemble means, and the average width and coverage of the 90% prediction intervals for NEE and LE after LOO-CV calibrations. While the joint calibration predictive intervals were narrower on average (1.3e-07 vs 1.5e-07 for NEE, 88 vs 103 for LE), the percentage of observations that fall into the predictive intervals were lower compared to the hierarchical calibration (80% vs 84% for NEE, 73% vs 76% for LE). The performance metrics of MAP runs and ensemble means were very similar on average for both calibrations. Boxplots for deviance values of each ensemble member are given in Fig S6 which shows that while joint calibration LOO-CV deviance values were lower on average, the best ensemble member (== lowest deviance) was usually in HB calibration LOO-CV ensemble.

**Fig 7.**
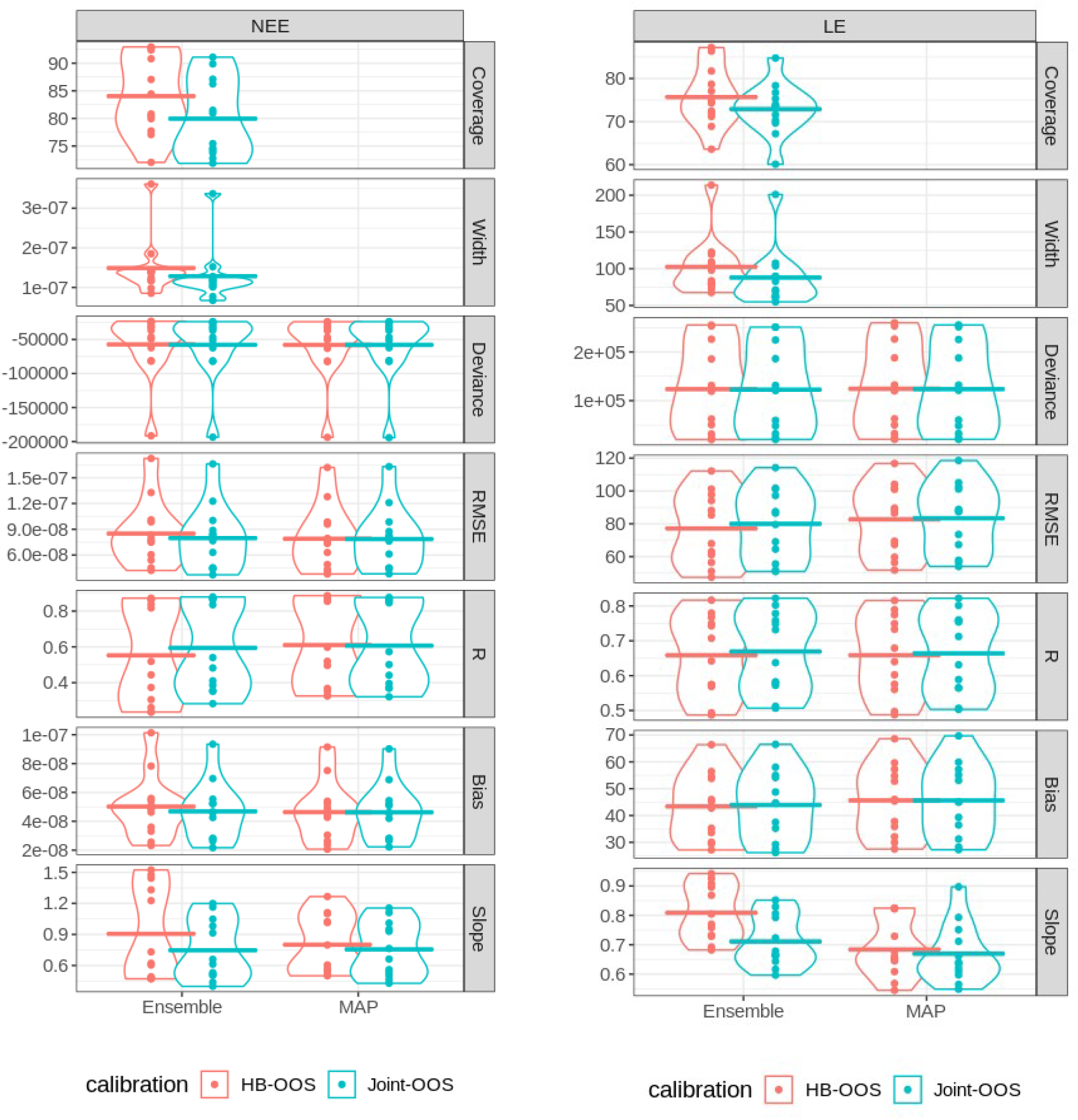
Performance metrics comparing HB versus Joint calibration at the out-of-sample (OOS) sites. Coverage and width are calculated for the %90 predictive interval of the post-calibration ensembles. Rest of the metrics are provided for both ensemble mean and the maximum a posteriori (MAP) run. Each point is a site.

### 3.5 Uncertainty partitioning

(H4) Across-site parameter variability will be higher than parameter uncertainty.

Partitioning of effects of the HB calibration by propagating the parameter uncertainty encapsulated by the site-level parameter posterior distributions, and the parameter variability encapsulated by the global parameter posterior distributions into model predictions revealed that uncertainty is dominated by across-site parameter variability (Fig 7), confirming our 4^th^ hypothesis.

**Fig 7.**
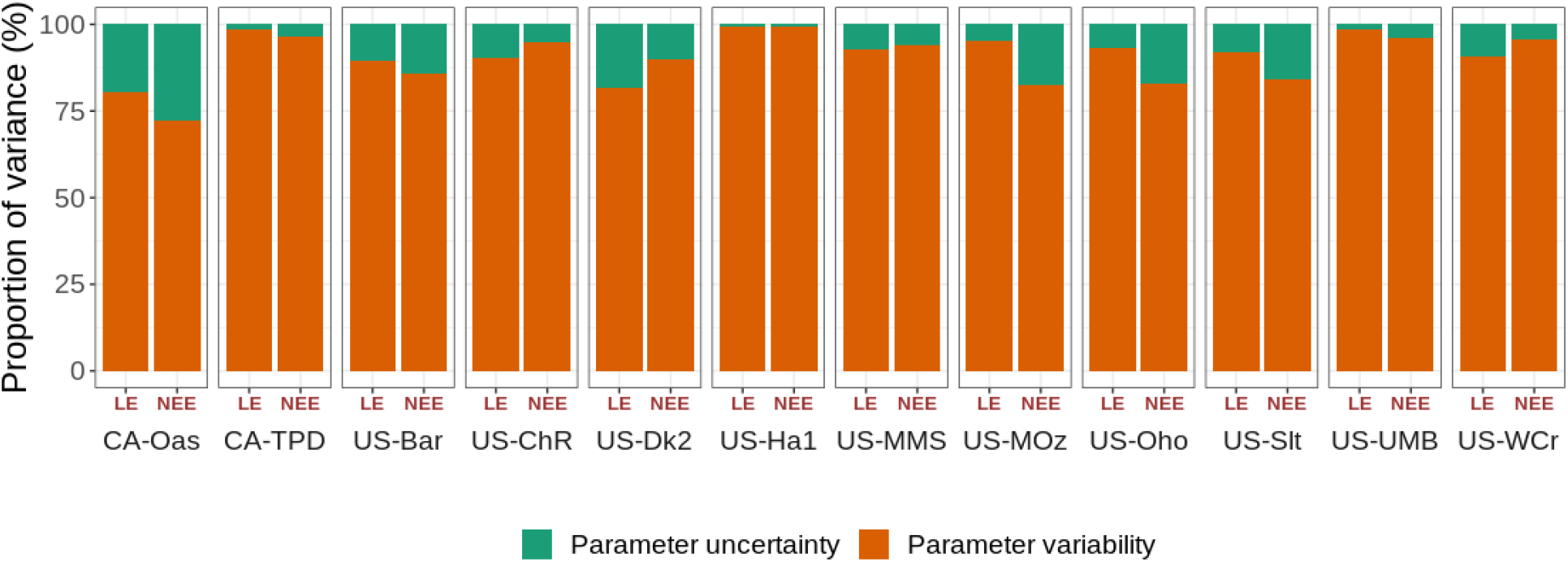
Partitioning of model output uncertainties by parameter variability versus parameter uncertainty at the sites.

## 4. Discussion

### 4.1 In-sample experiments

Comparison of individual versus joint versus hierarchical Bayesian calibration in this study supported our prior theoretical expectations. Site-level calibrations under SB and HB calibration showed the best agreement with data while joint calibration, where all data from all sites were used in the calibration simultaneously to estimate a single parameter vector, returned the narrowest posterior parameter distributions and the narrowest confidence intervals for post-calibration ensemble runs. However the goodness-of-fit (deviance) of joint calibration was worse than the other two approaches. With HB calibration, site-level posteriors were able to gain further constraint in terms of both reduction in posterior parameter uncertainty, post-calibration ensemble uncertainty and increase in goodness-of-fit. The differences in post-calibration performances of the three approaches were admittedly small; however, we note that this was a result from targeting five model parameters and using only twelve sites in this proof of concept study. With more parameters and more sites included in the calibration the false confidence and lack of flexibility of the joint calibration is expected to get worse, while the gains from HB calibration are expected to become more pronounced.

### 4.2 Analysis of the random effects

Analysis of across-site correlation showed relatively modest (negative) correlation between across-site parameters, the highest being between the aerodynamic resistance constant and the leaf growth parameters. In other words, sites with high aerodynamic resistance constants were having low leaf growth values, and vice versa. In SIPNET, both higher aerodynamic resistance and leaf growth values result in less evaporation from soil surface. However, except for one site, no similar trade-off was observed within sites (not shown), therefore it is unlikely that it could have led to across-site trade-offs. Still, such across-site correlations have practical implications. For example, if we can inform one of these correlated parameters when making out-of-sample predictions, we would gain information about the other (Shiklomanov et al., 2020).

The correlation analysis between potential explanatory variables and site-to-site parameter variability showed relatively high correlation between mean annual temperature and optimum photosynthesis parameter as expected (Dietze, 2014; Lombardozzi et al., 2015). This suggests that the model may be trying to compensate for the response to different temperatures by using different values for these parameters, and thus improving the temperature sensitivity, or possibly accounting for acclimation, ecotypic adaptation, or species composition within the model processes can reduce site-to-site variability and improve model predictions. Separating acclimation from adaptation is difficult from the current analysis, but could be improved by further analyses exploring different random effects such as temporal random effects if acclimation is over shorter timescales.

The highest site-to-site variability, however, was seen for the soil respiration parameter, and the highest correlations between explanatory variables and its site-to-site variability were with initial carbon pools. This could be a compensating error in SIPNET. It could also be due to missing processes in SIPNET where the model cannot capture that larger pools are associated with more recalcitrant carbon. This supports recent directions of improvements to use realistic and measurable carbon pools, and include coupled carbon, nitrogen and phosphorus cycles in soil biogeochemical modeling (Abramoff et al., 2018; Thum et al., 2019).

Among explanatory variables, a priori fixed parameters for the Laplace error model of fluxes also showed high correlations with site-to-site variability of some parameters, especially with the water use efficiency constant (wueConst). wueConst was the most influential parameter contributing to the model output uncertainties for the latent heat (LE) variable. In other words, Laplace parameters related to LE data stream may be expected to explain some of the site-to-site variability in wueConst. However, within site parameter correlations (not shown) also show relatively high negative correlation between wueConst and optimum photosynthesis parameter (psnTOpt) or positive correlation between wueConst and leaf growth parameter, sometimes both. Both psnTOpt and leaf growth parameter values were also controlled by the NEE data stream, therefore Laplace parameters related to NEE may also be expected to explain some of the site-to-site variability in wueConst within our current calibration scheme, which was the case for our results. However, in principle, the parameters of the error model should not be correlated with site-to-site variability of the process-model parameters. We calculated the parameters for the Laplace error model following the approach in Richardson et al. (2006). Fits for the asymmetric heteroskedastic Laplace flux uncertainty parameters of NEE (Fig S5) revealed that the US-Dk2 site has distinctly different slopes than the rest of the sites. Indeed, when this site was removed from the correlation analysis there was no remaining correlation relationship between fixed Laplace parameters and site-to-site variability of model parameters. A closer look at the raw flux data from this site showed noisy data even after the u* filtering which needs further inspection. Still, an apparent next step for our calibration scheme is to fit the Laplace error model parameters simultaneously with the rest of the model parameters, instead of using a priori fixed values.

### 4.2 Out-of-sample experiments

Leave-one-out cross validation (LOO-CV) exercise showed that even though joint calibration global posterior ensemble simulations were more confident (smaller width on average), they excluded more of the observational data points in their predictive intervals. This was also visible in the residual check (not shown) where joint calibration posterior predictive intervals over- and under-predicted more values in comparison to the HB LOO-CV. We note that even fewer number (11) of sites went into these LOO-CV calibrations which in return reflects the impact of different calibration schemes rather weakly. Regardless, the results supported our initial hypotheses partially where the joint calibration was overconfident, but they were not overconfidently-wrong as hypothesized. The model predictions using the HB calibration were not necessarily more accurate at the out-of-sample sites, but more honest about their true uncertainty. Nevertheless, HB global posteriors provide a formal prior for a site-level calibration at the out-of-sample sites. In other words, when new or more data becomes available from a site that did not go into the hierarchical calibration, we can perform an individual site-level calibration using previous analysis’ HB global parameters as prior. This would not be possible with joint calibration as its over-confident global posteriors would practically overwhelm the new information from individual sites.

Furthermore, propagation of parameter uncertainty and parameter variability to the model predictions suggests that parameter variability was more important than parameter uncertainty considering the five parameters targeted in this study. This result suggests that instead of making more or longer flux measurements in the existing sites, we need flux measurements at more sites. For example, measurement campaigns such as the Chequamegon Heterogeneous Ecosystem Energy-Balance Study Enabled by a High-Density Extensive Array of Detectors 2019 (CHEESEHEAD19, Butterworth et al., 2021) where an EC tower network was deployed to sample land-cover variation would be the most informative in this context.

### 4.3 Future steps

In this study we included 12 sites as we worked with a simple model that can simulate one biome type at a time. Next step is to perform emulator-based hierarchical Bayesian calibration of a more sophisticated dynamic vegetation model which would allow us to include more sites that can better inform the site effects. However, emulating such models might require emulator implementations that are more scalable because in that case there would be more parameters (of multiple plant functional types that represent different biomes) to target in the calibration. Adequately training an emulator for a parameter space of increasing size requires more design points, and our current emulator also gets slow with size due to O(N^3^) floating point operations and O(N^2^) memory that are needed in the Gaussian Process (GP) implementation, limiting its applicability in the current calibration scheme. Therefore, utilizing recent sparsity solutions to computational limitations of full GPs would be the logical next step in this regard (Vanhatalo et al., 2013; Matthews et al., 2017).

Furthermore, at the moment our hierarchical cross-site calibration treats site-to-site parameter differences as simple “random effects” that are independent and uncorrelated. However, as we switch to more scalable emulators and increase the number of calibration sites (with the distance between them decreasing) we need to better consider the spatial correlation structure in parameter variability. Similarly, our hierarchical model here only considered spatial variability in model parameters, but just as there are unaccounted for processes that cause variability in space, there are likely to be unaccounted for processes in models that would cause the parameters to vary in time (Wikle et al., 2019). Therefore a logical next step is to develop spatiotemporal models of parameter variability at the hierarchical level using our the emulator framework.

## Supplementary material

**Fig S1.**
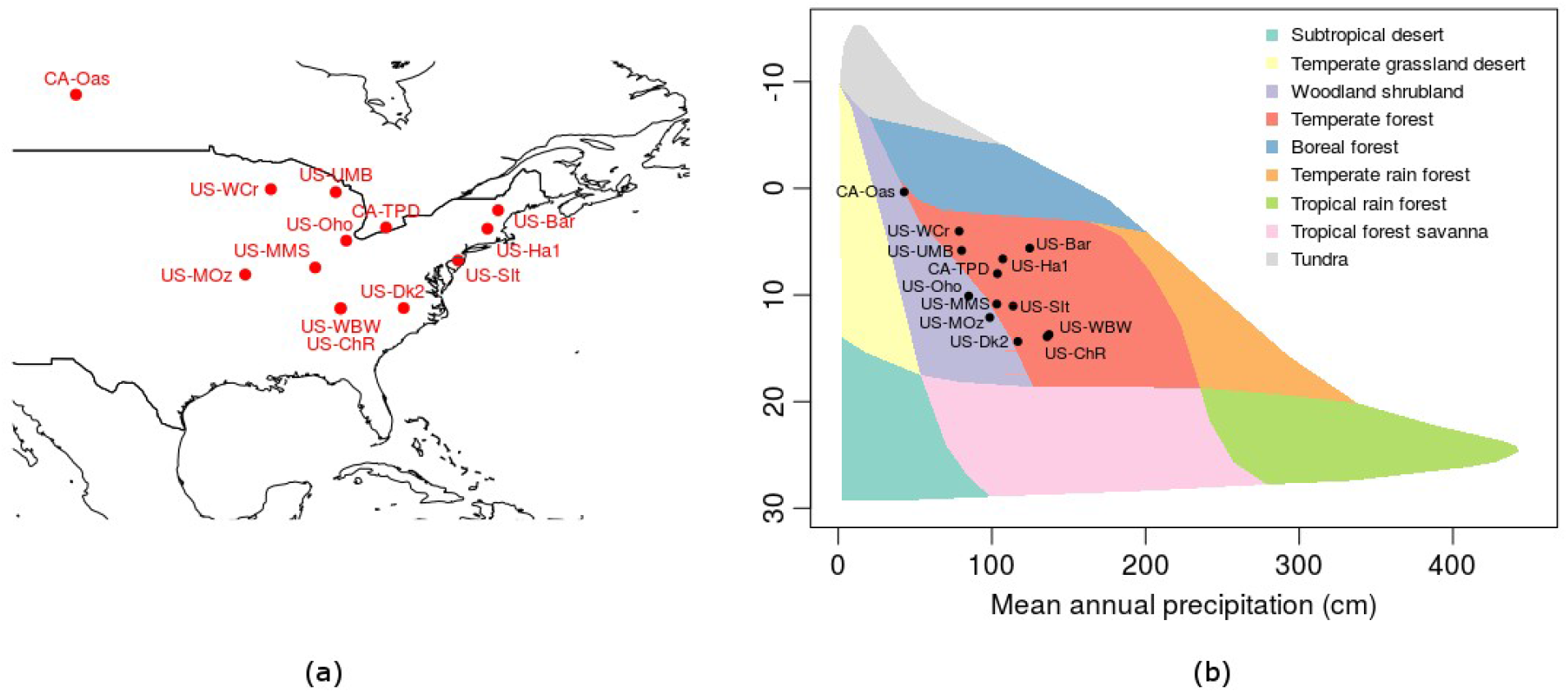
Sites in the geographic (a) and climatic (b) space.

**Fig. S2:**
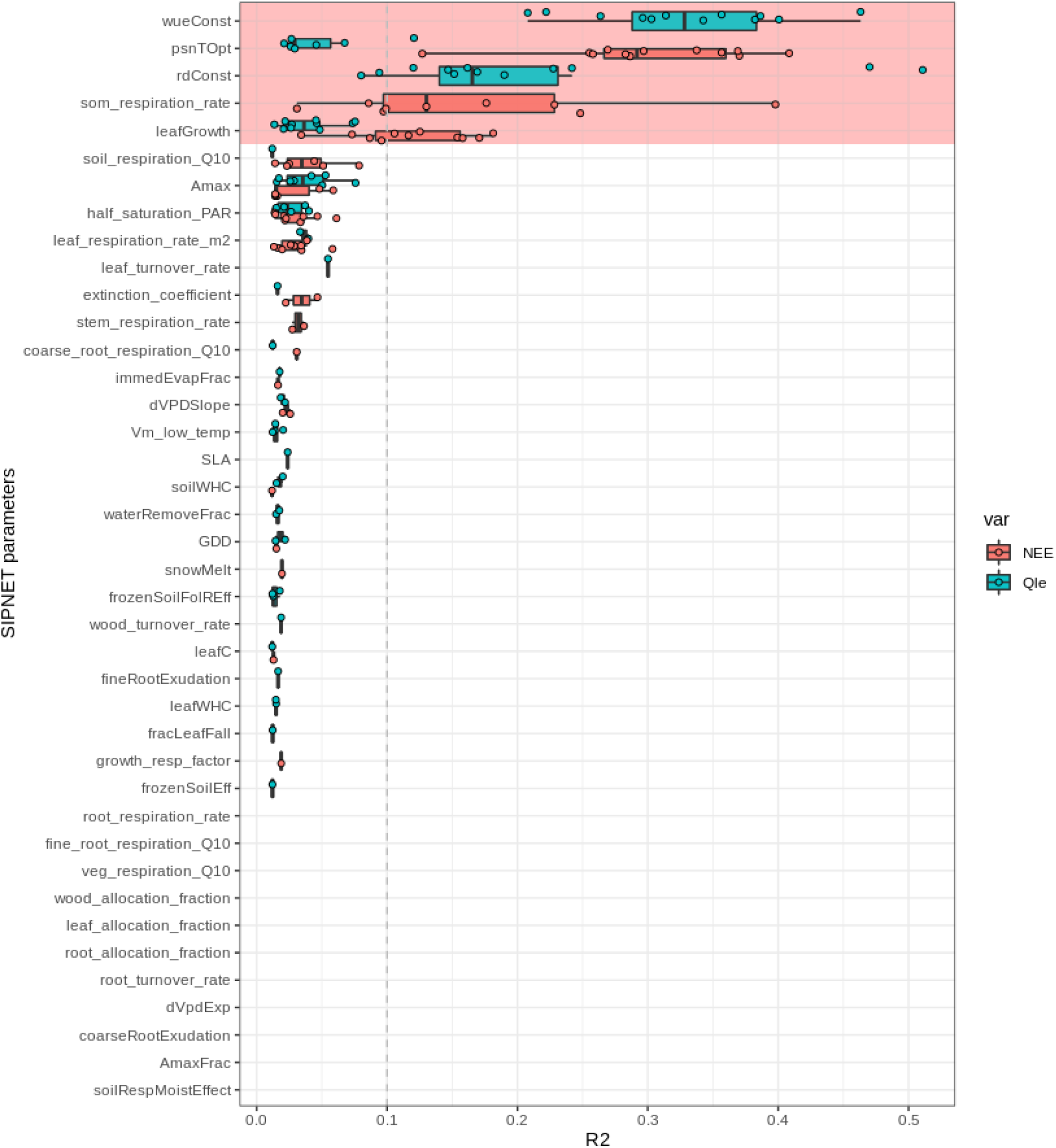
The results of the uncertainty analysis. Box and whisker plots show the relative importance of parameters in terms of R^2^ values from fitted linear regressions between each parameter sample and model outputs, the higher the R^2^ the more the parameter contributes to the output uncertainty. Each point represents a site for the parameters with significant fits (F-statistic, p-value < 0.05). Vertical dashed line represents the 10% threshold under which we considered the relative contribution of the parameters to the model output uncertainty negligible and omitted from the calibration. In the end, the top 5 SIPNET parameters in the shaded area were chosen to be targeted in the calibration.

**Table S1.** Ameriflux Deciduous Broadleaf Forest sites.

| Site ID | Site name | Lat | Lon | Years used | # years |
| --- | --- | --- | --- | --- | --- |
| US-Bar | Bartlett Experimental Forest | 44.0646 | -71.2881 | 2005-2011 | 7 |
| US-ChR | Chestnut Ridge | 35.9311 | -84.3324 | 2008-2010 | 3 |
| US-Ha1 | Harvard Forest EMS Tower | 42.5378 | -72.1715 | 2010-2015 | 6 |
| US-Dk2 | Duke Forest hardwoods | 35.9736 | -79.1004 | 2002-2008 | 7 |
| US-MMS | Morgan Monroe State Forest | 39.3232 | -86.4131 | 2001-2014 | 14 |
| US-MOz | Missouri Ozark Site | 38.7441 | -92.2 | 2005-2015 | 11 |
| US-Oho | Oak Openings | 41.5545 | -83.8438 | 2004-2013 | 10 |
| US-Slt | Silas Little | 39.9138 | -74.596 | 2005-2014 | 10 |
| US-UMB | Univ. of Mich. Biological Station | 45.5598 | -84.7138 | 2007-2014 | 8 |
| US-WCr | Willow Creek | 45.8059 | -90.0799 | 1999-2006 | 8 |
| CA-TPD | Turkey Point Mature Deciduous | 42.6353 | -80.5577 | 2013-2015 | 3 |
| CA-Oas | BOREAS SSA Old Aspen | 53.6289 | -106.1978 | 1997-2010 | 14 |

**Table S2.** SIPNET initial conditions.

| Symbol | SIPNET Code | Definition | Units |
| --- | --- | --- | --- |
| $C_W$ | PlantWoodInit | Initial plant wood C content ($C_A$:above ground + $C_R$:roots) | gC m<sup>-2</sup> |
| $LAI_0$ | lailnit | Initial leaf area index | m<sup>2</sup> m<sup>-2</sup> |
| $C_L$ | litterInit | Initial litter C content ($C_{LL}$:leaf + $C_{LW}$:woody) | gC m<sup>-2</sup> |
| $C_S$ | soilInit | Initial soil C content | gC m<sup>-2</sup> |
| $W_{SP}$ | snowInit | Initial snow pack | cm water equivalent |

**Table S3.** Initial pool values. LAI0 are set to 0 for all sites, as runs start from winter dormancy. W_SP_ are also set to 0 for all sites following previous studies, as this is a fast pool and snow accumulation by Jan 1^st^ is assumed to be negligible at most of our sites. When only one of them or the total wood C content were available in the literature, 0.8/0.2 ratio is assumed for C_A_/C_R_. When only one of them or the total litter C content were available in the literature, 0.5/0.5 ratio is assumed for C_LL_/C_LW_.

| Site | $C_A$ | $C_R$ | $C_{LL}$ | $C_{LW}$ | $C_S$ |
| --- | --- | --- | --- | --- | --- |
| US-Bar | 10600 | 2000 | 110 | 65 | 1600 |
| US-ChR | 7220 | 1910 | 170 | 170 | 5570 |
| US-Ha1 | 12125 | 3000 | 219 | 219 | 3943 |
| US-Dk2 | 18000 | 2900 | 216 | 250 | 4360 |
| US-MMS | 10588 | 2500 | 190 | 190 | 6600 |
| US-MOz | 10700 | 2500 | 232 | 232 | 1800 |
| US-Oho | 9950 | 2000 | 194 | 194 | 8225 |
| US-Slt | 5000 | 1250 | 198 | 198 | 1800 |
| US-UMB | 10120 | 2500 | 148 | 168 | 5189 |
| US-WCr | 7100 | 1800 | 182 | 182 | 5350 |
| CA-TPD | 12300 | 3000 | 193 | 193 | 5500 |
| CA-Oas | 7550 | 980 | 160 | 160 | 5533 |

**Table S4.** Compilation sources for initial conditions. If data for multiple years are provided in the source, the closest year to the year before the start date was used.

| Site ID | IC Pool | Source | Source info |
| --- | --- | --- | --- |
| US- Bar | $C_A$ | BADM file | AG_BIOMASS_TREE_ORGAN (Wood) |
| | $C_R$ | BADM file | ROOT_BIOMASS |
| | $C_{LL}$ | BADM file | AG_LIT_BIOMASS (leaf litter) |
| | $C_{LW}$ | BADM file | AG_LIT_BIOMASS (large branches+other) |
| | $C_S$ | Bradford et al., 2010 | Table 2 |
| US-ChR | C<sub>A</sub> | Malhi et al. 1999 | Approximated from Malhi et al. 1999, Table 5 |
|  | C<sub>R</sub> | Malhi et al. 1999 | Approximated from Malhi et al. 1999, Table 5 |
|  | C<sub>LL</sub> | Davidson et al. 2002 | Approximated from Davidson et al. 2002 |
|  | C<sub>LW</sub> | Davidson et al. 2002 | Approximated from Davidson et al. 2002 |
|  | C<sub>S</sub> | Malhi et al. 1999 | Approximated from Malhi et al. 1999, Table 5 |
| US-Ha1 | C<sub>A</sub> | BADM file | AG_BIOMASS_TREE_ORGAN (Wood) |
|  | C<sub>R</sub> | BADM file | Approximated from C<sub>A</sub> |
|  | C<sub>LL</sub> | Davidson et al. 2002 | Approximated from Davidson et al. 2002 |
|  | C<sub>LW</sub> | Davidson et al. 2002 | Approximated from Davidson et al. 2002 |
|  | C<sub>S</sub> | BADM file | SOIL_STOCK_C_ORG |
| US-Dk2 | C<sub>A</sub> | BADM file | AG_BIOMASS_TREE_ORGAN (Wood) |
|  | C<sub>R</sub> | BADM file | ROOT_BIOMASS |
|  | C<sub>LL</sub> | BADM file | AG_LIT_BIOMASS |
|  | C<sub>LW</sub> | BADM file | WD_BIOMASS_CRS |
|  | C<sub>S</sub> | BADM file | SOIL_CHEM_C_ORG + SOIL_CHEM_BD |
| US-MMS | C<sub>A</sub> | BADM file | AG_BIOMASS_TREE_ORGAN (Wood) |
|  | C<sub>R</sub> | BADM file | Approximated from C<sub>A</sub> |
|  | C<sub>LL</sub> | BADM file | AG_LIT_BIOMASS |
|  | C<sub>LW</sub> | BADM file | Approximated from C<sub>LL</sub> |
|  | C<sub>S</sub> | BADM file | SOIL_STOCK_C_ORG |
| US-MOz | C<sub>A</sub> | Hanberry et al 2016 | Approximated value |
|  | C<sub>R</sub> | Hanberry et al 2016 | Approximated value |
|  | C<sub>LL</sub> | BADM file | AG_LIT_BIOMASS (total) |
|  | C<sub>LW</sub> | BADM file | AG_LIT_BIOMASS (total) |
|  | C<sub>S</sub> | Hammer 1997 | Approximated from soil C content |
| US-Oho | C<sub>A</sub> | Noormets et al. 2014 | Study site description |
|  | C<sub>R</sub> | Noormets et al. 2014 | Study site description |
|  | C<sub>LL</sub> | Noormets et al. 2014 | Study site description |
|  | C<sub>LW</sub> | Noormets et al. 2014 | Study site description |
|  | C<sub>S</sub> | Noormets et al. 2014 | Study site description |
| US-Slt | C<sub>A</sub> | BADM file | AG_BIOMASS_ORGAN (Wood) |
|  | C<sub>R</sub> | BADM file | Approximated from C<sub>A</sub> |
|  | C<sub>LL</sub> | BADM file | AG_LIT_BIOMASS (total) |
|  | C<sub>LW</sub> | BADM file | AG_LIT_BIOMASS (total) |
|  | C<sub>S</sub> | BADM file | SOIL_CHEM_C_ORG + SOIL_TEX |
| US-UMB | C<sub>A</sub> | BADM file | AG_BIOMASS_TREE_ORGAN (Wood) |
|  | C<sub>R</sub> | BADM file | ROOT_BIOMASS_TOT |
|  | C<sub>LL</sub> | Davidson et al. 2002 | Approximated from Davidson et al. 2002 |
|  | C<sub>LW</sub> | Davidson et al. 2002 | Approximated from Davidson et al. 2002 |
|  | C<sub>S</sub> | BADM file | SOIL_CHEM_C_ORG + SOIL_CHEM_BD |
| US-WCr | C<sub>A</sub> | Site PI | Personal communication |
|  | C<sub>R</sub> | Site PI | Personal communication |
|  | C<sub>LL</sub> | Bolstad et al. 2004 | Aboveground litter input |
|  | C<sub>LW</sub> | Bolstad et al. 2004 | Approximated from C<sub>LL</sub> |
|  | C<sub>S</sub> | Cook et al. 2004 | Site description and flux source area |
| CA-TPD | C<sub>A</sub> | Site PI | Personal communication |
|  | C<sub>R</sub> |  | Approximated from C<sub>A</sub> |
|  | C<sub>LL</sub> | Site PI | Personal communication |
|  | C<sub>LW</sub> |  | Approximated from C<sub>LL</sub> |
|  | C<sub>S</sub> | Literature | Approximate value from Puric-Mladenovic et al 2016 |
| CA-Oas | C<sub>A</sub> | BADM file | AG_BIOMASS_TREE_ORGAN (Wood) |
|  | C<sub>R</sub> | BADM file | ROOT_BIOMASS_TOT |
|  | C<sub>LL</sub> |  |  |
|  | C<sub>LW</sub> |  |  |
|  | C<sub>S</sub> | BADM file | SOIL_CHEM_C_ORG + SOIL_CHEM_BD |

**Table S5.** Environmental variables (besides carbon pools on Table S3) used in the correlation analysis at the DBF sites.

| Site ID | MAT<br>(°C) | MAP | PAR | VP | WS | CH<br>(m) | Stand<br>Age (yrs) | LAI | FT |
| --- | --- | --- | --- | --- | --- | --- | --- | --- | --- |
| US-Bar | 7.6 | 1338 | 9117 | 968 | 1.3 | 20 | 100 | 4.7 | MB |
| US-ChR | 14.6 | 1331 | 11125 | 1293 | 1.0 | 19 | 100 | 4.8 | OH |
| US-Ha1 | 8.7 | 1191 | 9916 | 1027 | 2.0 | 23 | 81 | 3.4 | OH |
| US-Dk2 | 15.3 | 1069 | 10589 | 1427 | 2.1 | 25 | 90 | 7.0 | OH |
| US-MMS | 12.5 | 1078 | 6538 | 1244 | 3.5 | 27 | 70 | 4.8 | MIX |
| US-MOz | 13.4 | 924 | 7402 | 1256 | 2.9 | 24 | 77 | 4.0 | OH |
| US-Oho | 10.5 | 809 | 7060 | 1106 | 2.1 | 24 | 46 | 4.0 | Oak |
| US-Slt | 12.4 | 1065 | 7061 | 1178 | 1.9 | 9.5 | 90 | 4.8 | OH |
| US-UMB | 7.2 | 619 | 5336 | 895 | 3.6 | 23 | 92 | 3.8 | MB |
| US-WCr | 5.8 | 752 | 9589 | 969 | 2.7 | 24 | 70 | 5.4 | MB |
| CA-TPD | 8.8 | 920 | 8750 | 1074 | 3.3 | 26 | 90 | 4.3 | WO |
| CA-Oas | 2.0 | 479 | 8583 | 624 | 3.0 | 20 | 85 | 3.4 | Aspen |

MAT: mean annual temperature, MAP: mean annual precipitation, PAR: photosynthetically active radiation, VP: Vapor pressure, CH: Canopy height, WS: wind speed; FT: forest type (MB: Maple-Beech, OH: Oak-Hickory, WO: White Oak)

**Fig S3.**
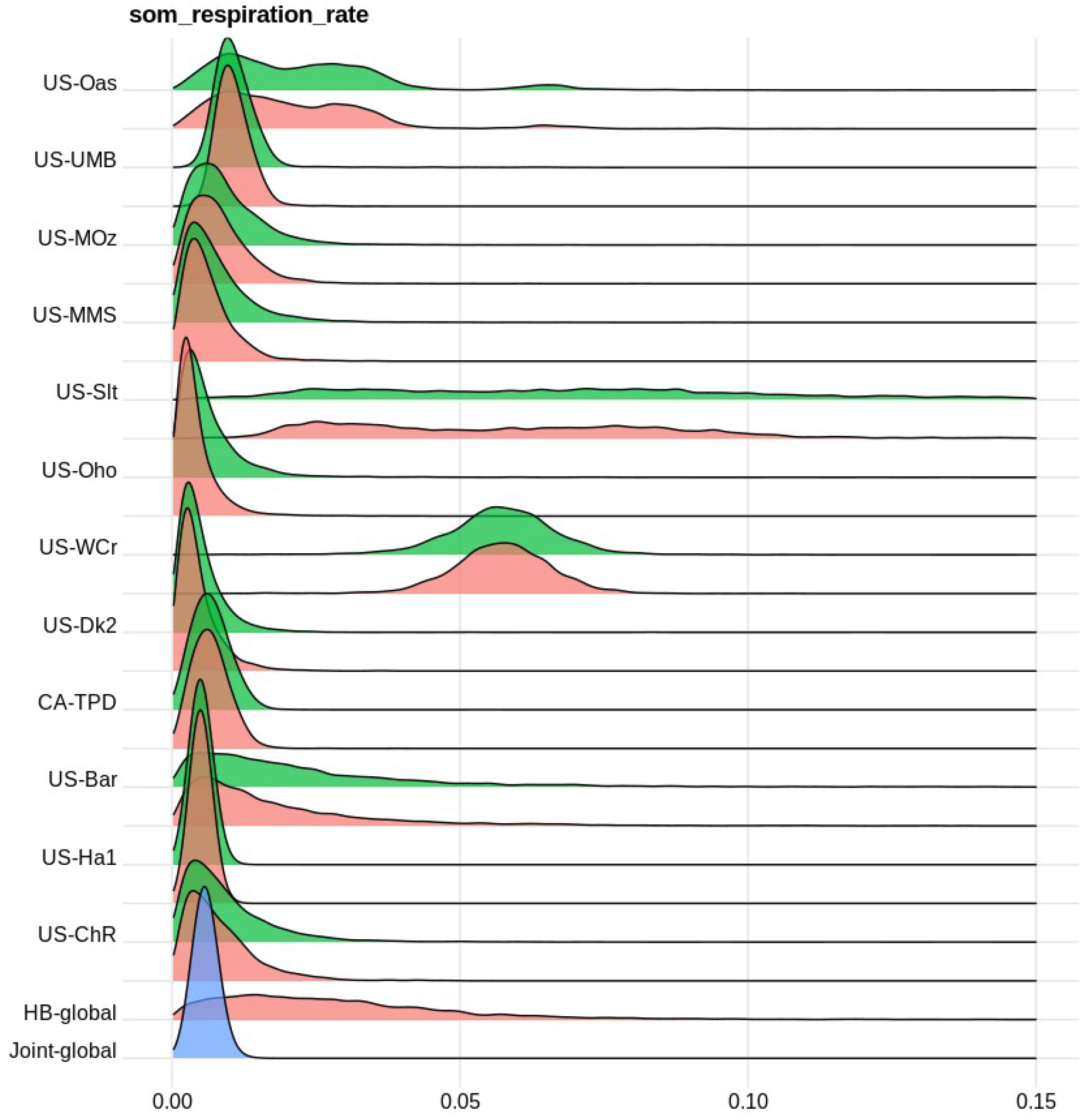
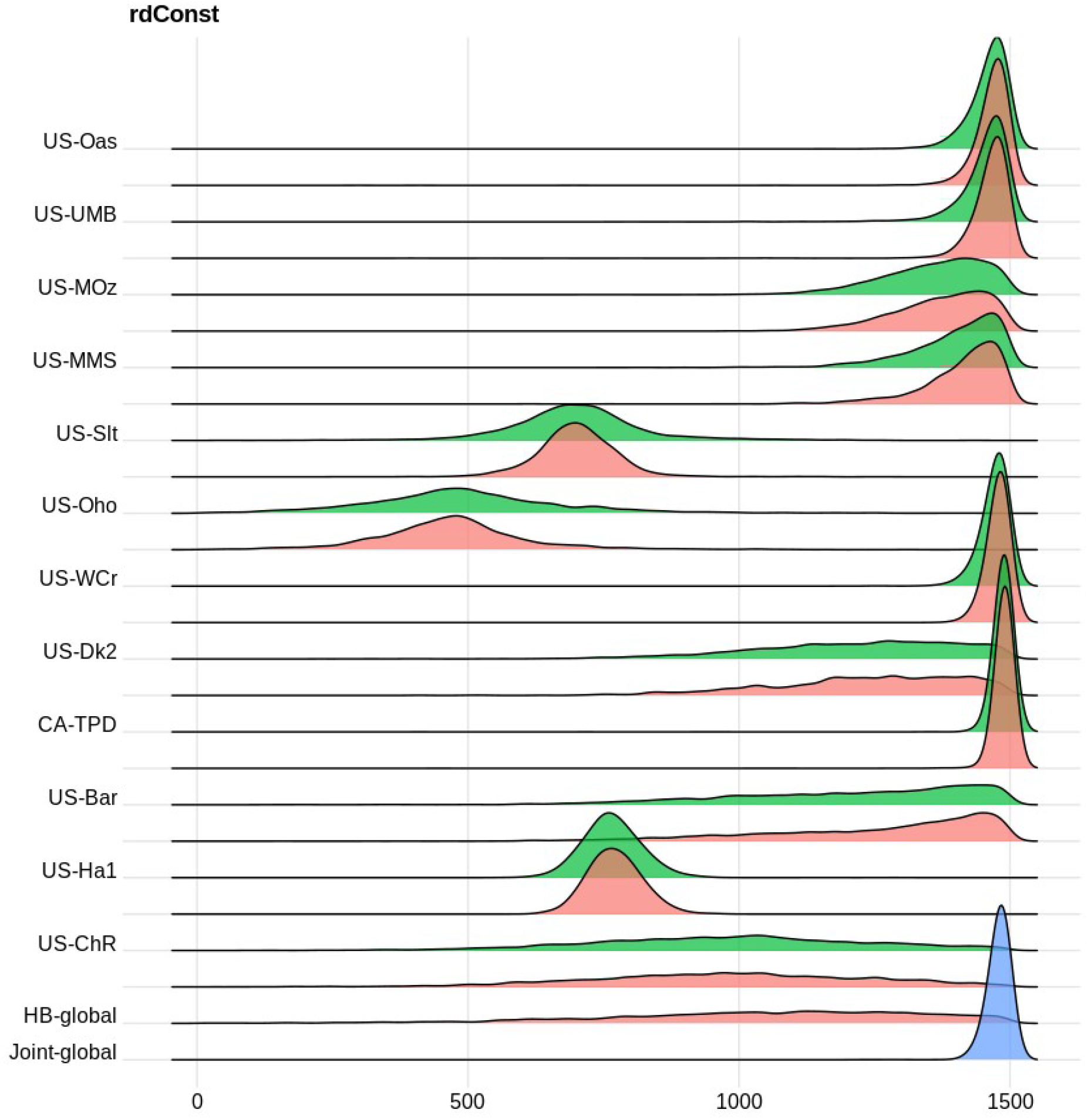
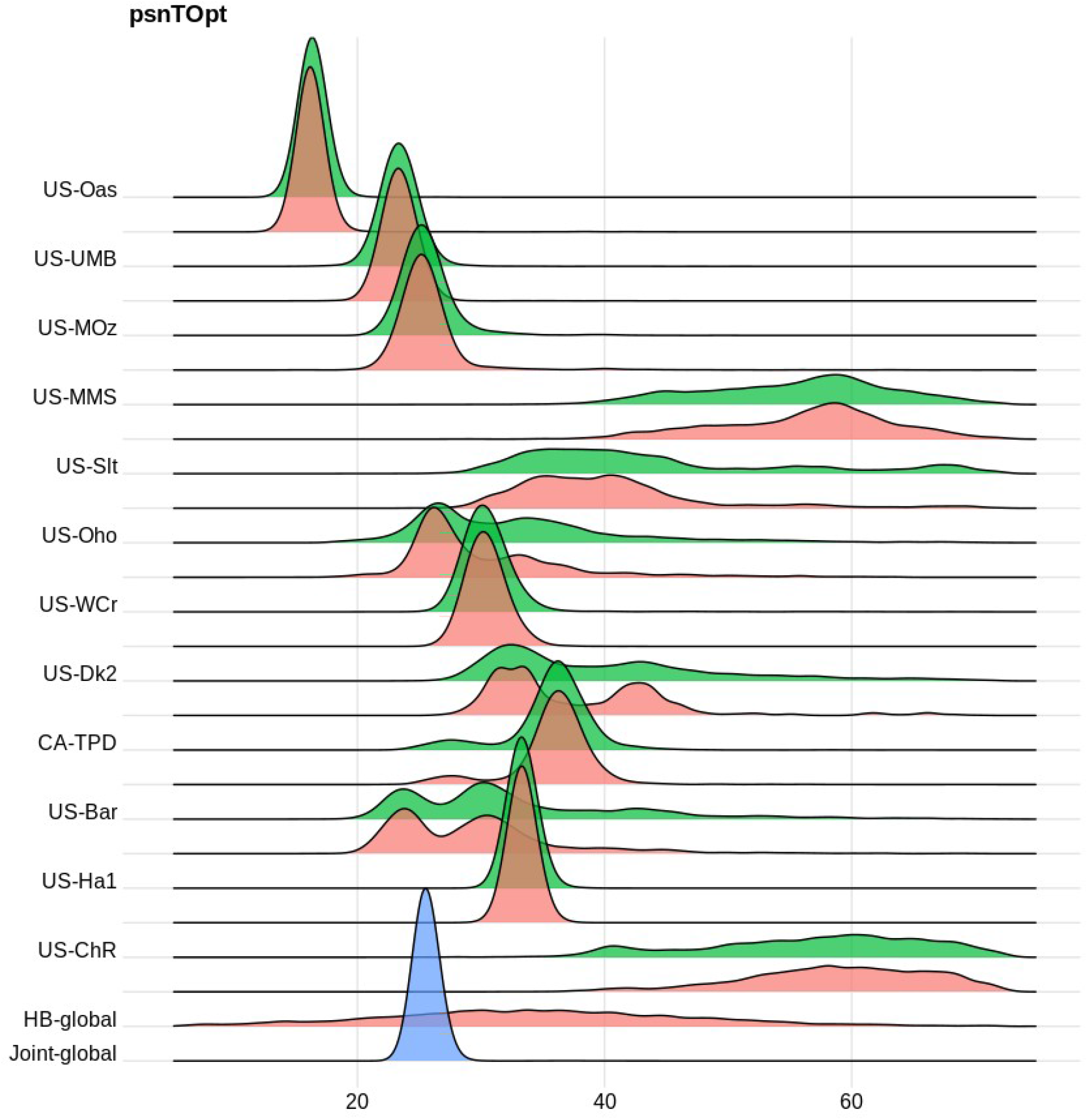
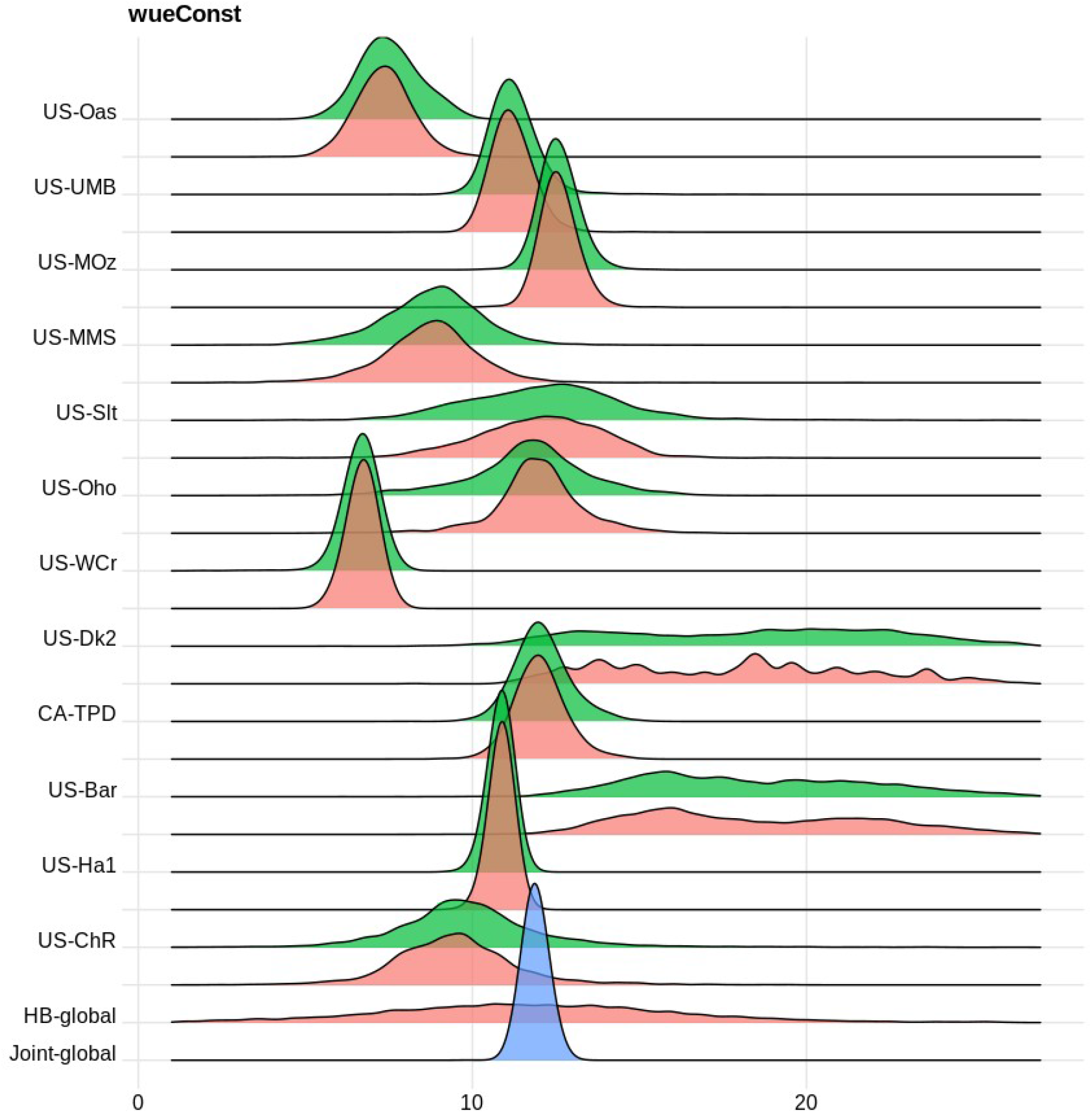
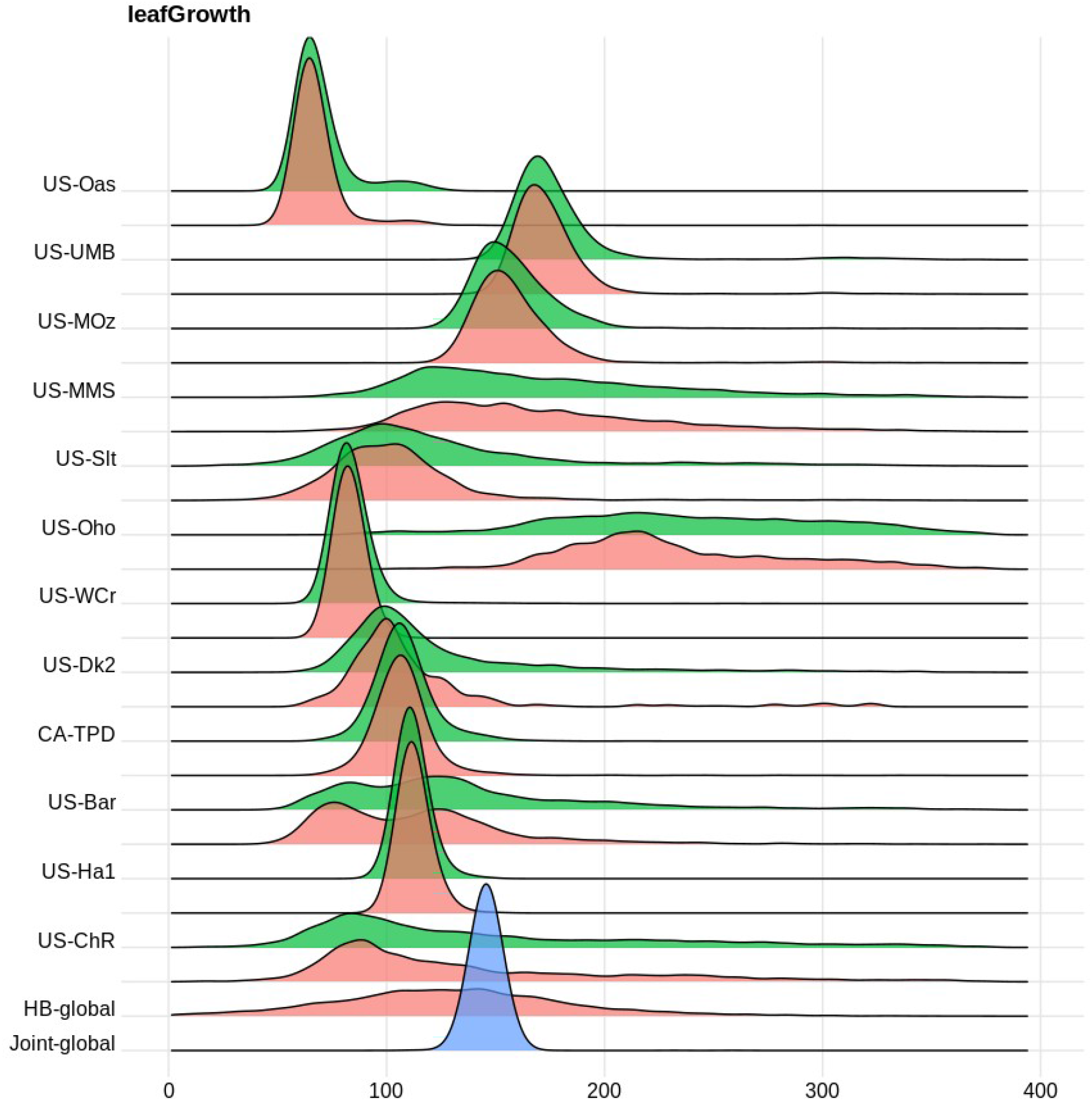
Marginal posterior distributions from each calibration approach for som_respiration_rate parameter. Joint calibration posterior (blue), SB site-level calibration posteriors (green) and hierarchical calibration posteriors (red). Cont’d for rdConst. Joint calibration posterior (blue), SB site-level calibration posteriors (green) and hierarchical calibration posteriors (red) Cont’d for psnTOpt. Joint calibration posterior (blue), SB site-level calibration posteriors (green) and hierarchical calibration posteriors (red) Cont’d for wueConst. Joint calibration posterior (blue), SB site-level calibration posteriors (green) and hierarchical calibration posteriors (red) Cont’d for leafGrowth. Joint calibration posterior (blue), SB site-level calibration posteriors (green) and hierarchical calibration posteriors (red)

**Fig S4.**
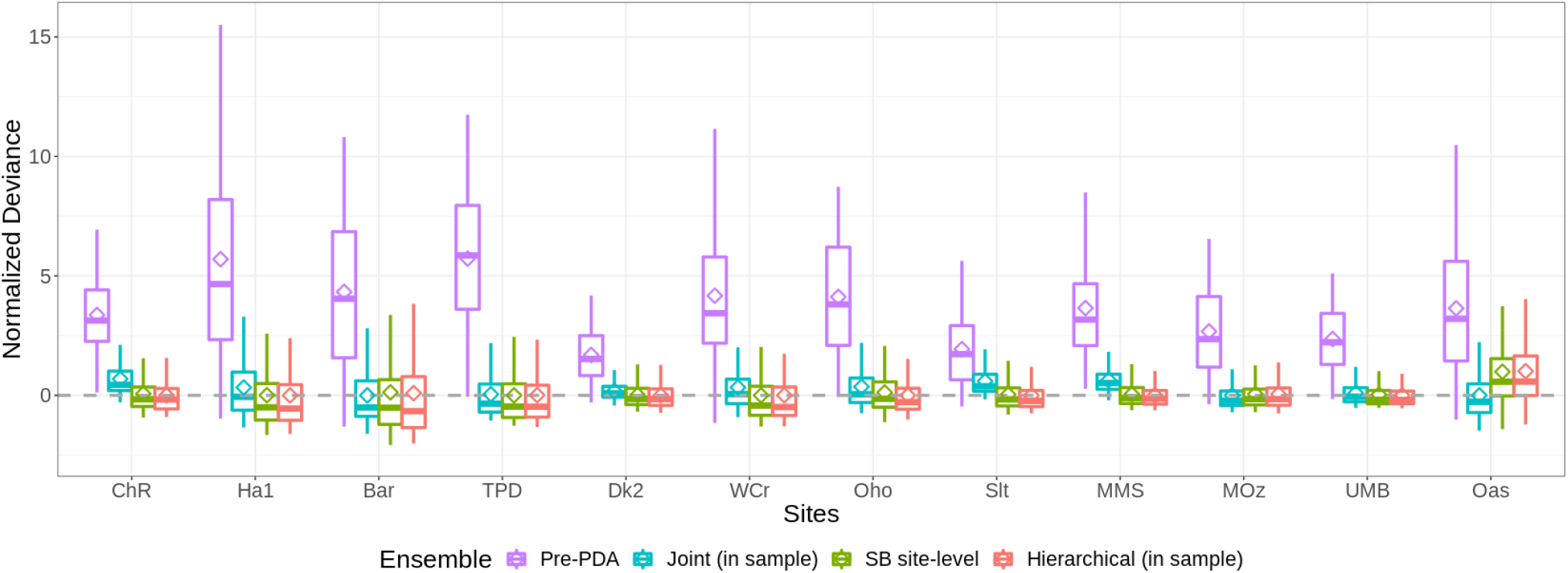
Same as Fig 6 but with pre-PDA. Normalized deviance results for ensemble runs (ensemble size 250) with different calibration approaches listed in Table 2. Deviance values are normalized by sample size at each site to obtain intercomparable results across sites, the lower the better.

**Fig S5.**
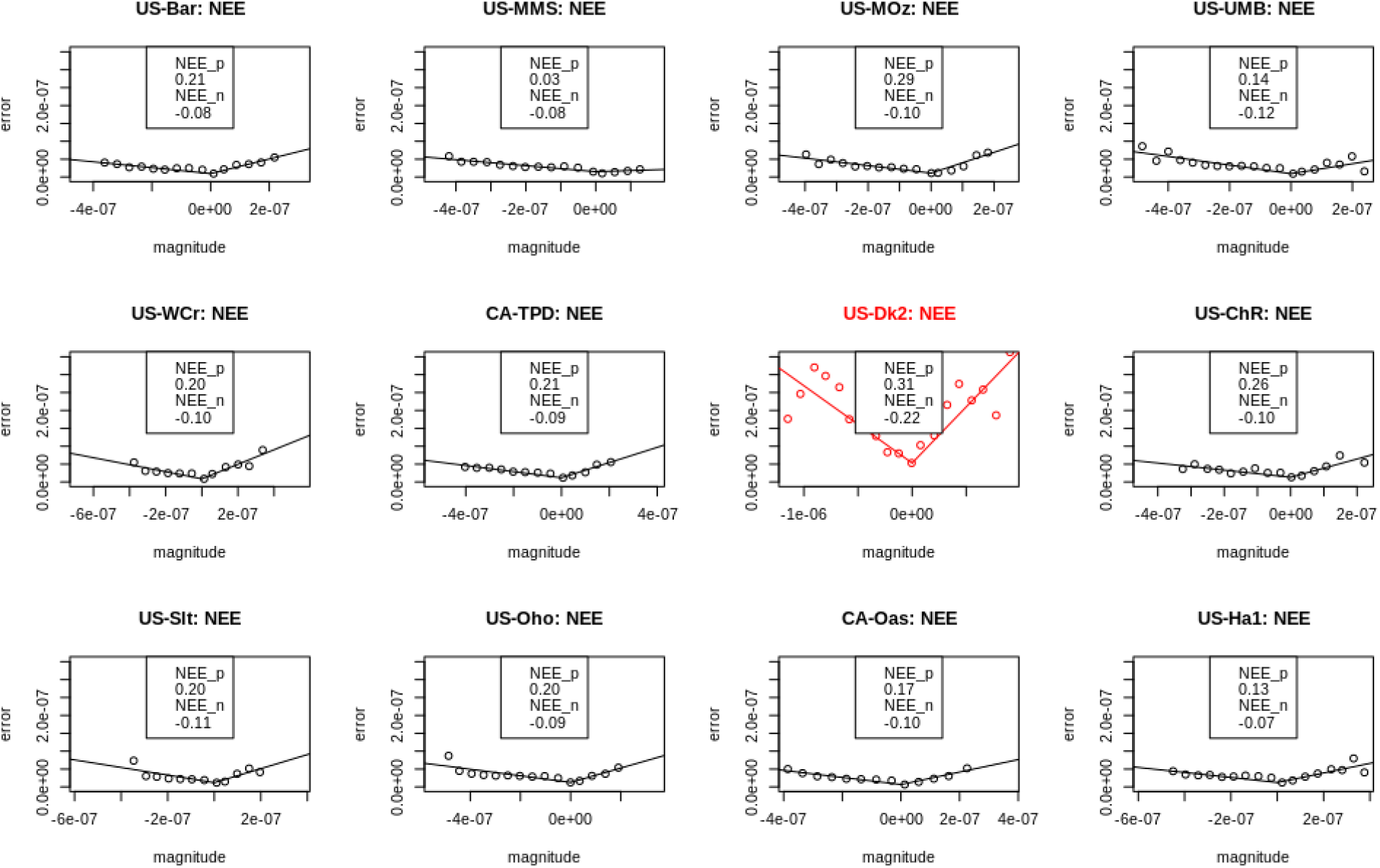
Fits for fixed a priori asymmetric heteroskedastic Laplace flux uncertainty parameters.

**Fig S6.**
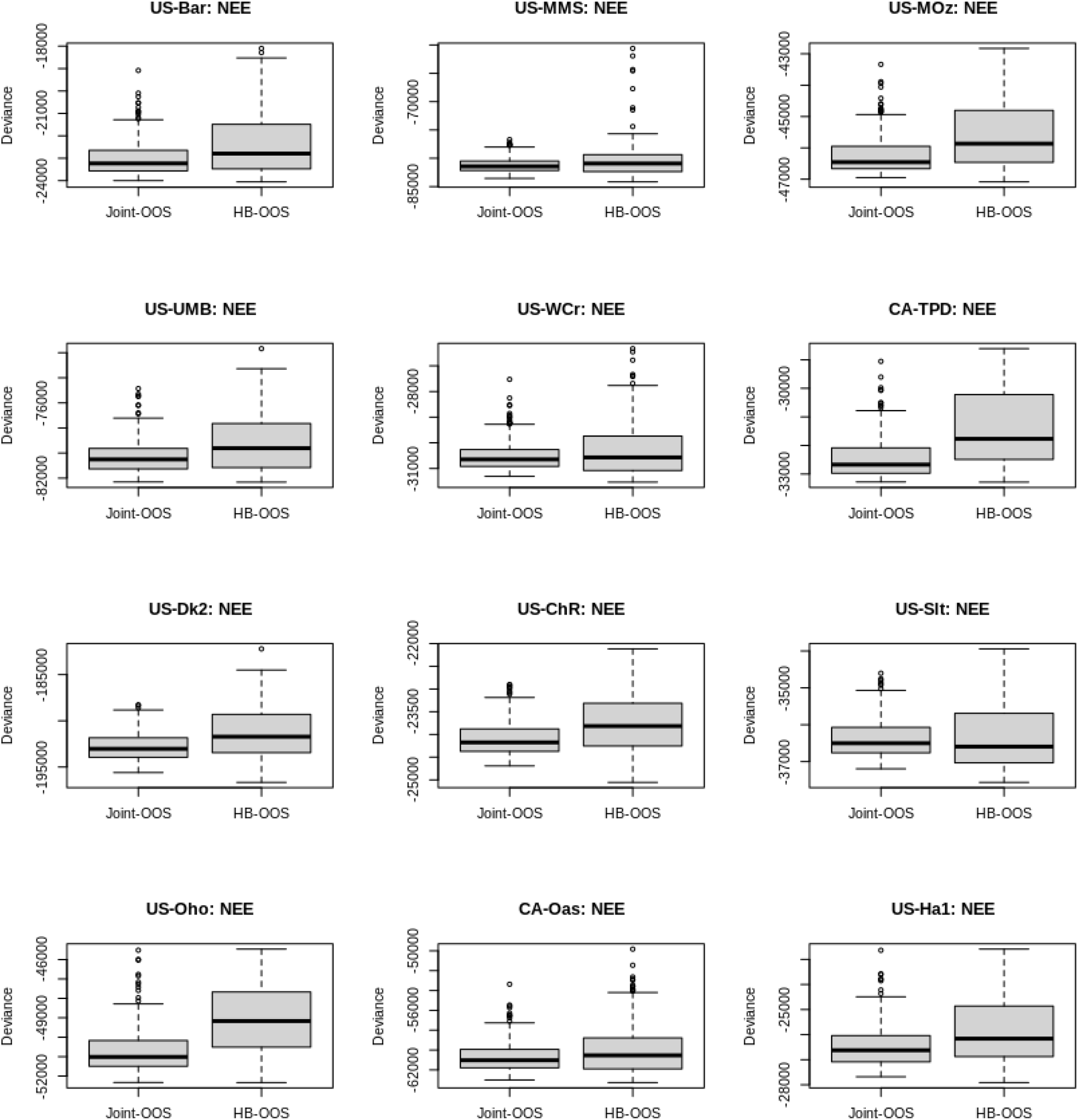
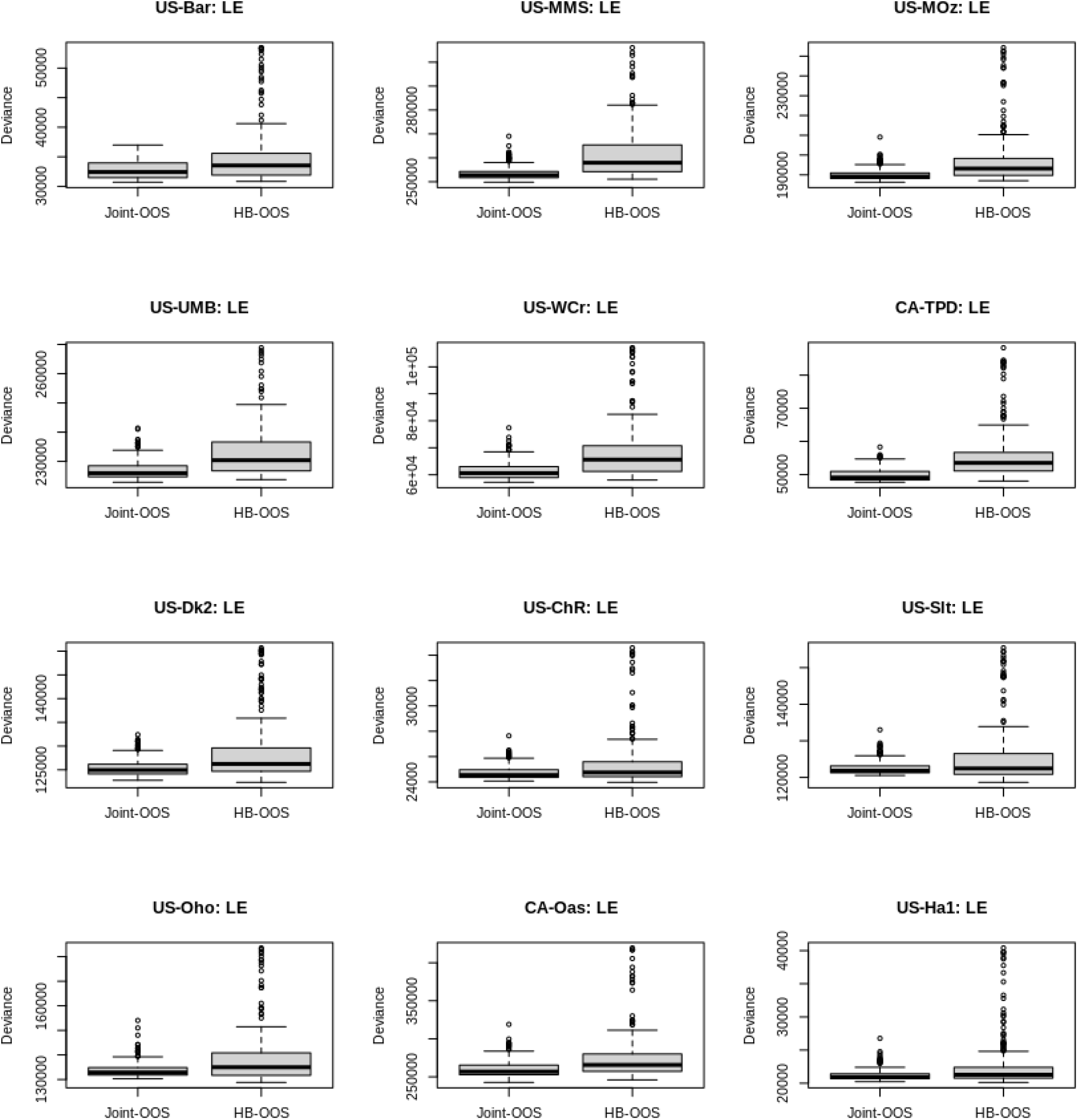
Deviance values of the ensemble members for NEE comparing HB versus Joint calibration at the out-of-sample (OOS) sites. Each boxplot consists of 250 ensemble members. cont’d for LE.

